# Spinophilin limits GluN2B-containing NMDAR activity and sequelae associated with excessive hippocampal NMDAR function

**DOI:** 10.1101/2020.12.30.424812

**Authors:** Asma B. Salek, Ruchi Bansal, Nicolas F. Berbari, Anthony J. Baucum

## Abstract

N-methyl-D-Aspartate receptors (NMDARs) are calcium-permeable ion channels that are ubiquitously expressed within the glutamatergic postsynaptic density. Phosphorylation of NMDAR subunits defines receptor activity and surface localization. Modulation of NMDAR phosphorylation by kinases and phosphatases regulates calcium entering the cell and subsequent activation of calcium-dependent processes. Spinophilin is the major synaptic protein phosphatase 1 (PP1) targeting protein that controls phosphorylation of myriad substrates via targeting or inhibition of PP1. Spinophilin limits NMDAR function in a PP1-dependent manner and we have previously shown that spinophilin sequesters PP1 away from the GluN2B subunit of the NMDAR, which results in increased phosphorylation of Ser-1284. However, how spinophilin modifies NMDAR function is unclear. Herein, we detail that while Ser-1284 phosphorylation increases calcium influx via GluN2B-containing NMDARs, overexpression of spinophilin decreases GluN2B-containing NMDAR activity by decreasing its surface expression. In hippocampal neurons isolated from spinophilin knockout animals there is an increase in cleaved caspase-3 levels compared to wildtype mice; however, this effect is not exclusively due to NMDAR activation; suggesting multiple putative mechanisms by which spinophilin may modulate caspase cleavage. Behaviorally, our data suggest that spinophilin knockout mice have deficits in spatial cognitive flexibility, a behavior associated GluN2B function within the hippocampus. Taken together, our data demonstrate a unique mechanism by which spinophilin modulates GluN2B containing NMDAR phosphorylation, channel function, and trafficking and that loss of spinophilin promotes pathological sequelae associated with GluN2B dysfunction.

**HIGHLIGHTS:**

- Spinophilin bidirectionally regulates GluN2B-containing NMDAR function.
- Loss of spinophilin in primary hippocampal neurons increases a pro-apoptotic marker.
- Loss of spinophilin *in vivo* decreases measures of spatial cognitive flexibility.

**Graphical Abstract:** Spinophilin increases the phosphorylation of Ser-1284 on GluN2B, thereby enhancing calcium influx through the GluN2B containing NMDARs. In contrast, spinophilin limits GluN2B-containing surface expression putatively due to modulation of GluN2B interactions with endocytotic proteins. Since the second effect of spinophilin occurs independent of the first, we observe an overall decrease in calcium influx through GluN2B containing NMDARs when spinophilin is present. This low, basal calcium influx is less likely to be promote calcium-dependent activation of caspase and downstream apoptotic pathways and permits flexible search strategies and behaviors. In the absence of spinophilin, the spinophilin-driven internalization of the receptors is decreased, more receptors are expressed on the surface and calcium influx into the cell is increased. This high levels of intracellular calcium triggers apoptotic pathways leading to cell death. This impact may be more dramatic in cells with high expression of GluN2B-containing NMDA receptors. This loss of spinophilin reduces cognitive flexibility in hippocampal dependent tasks.

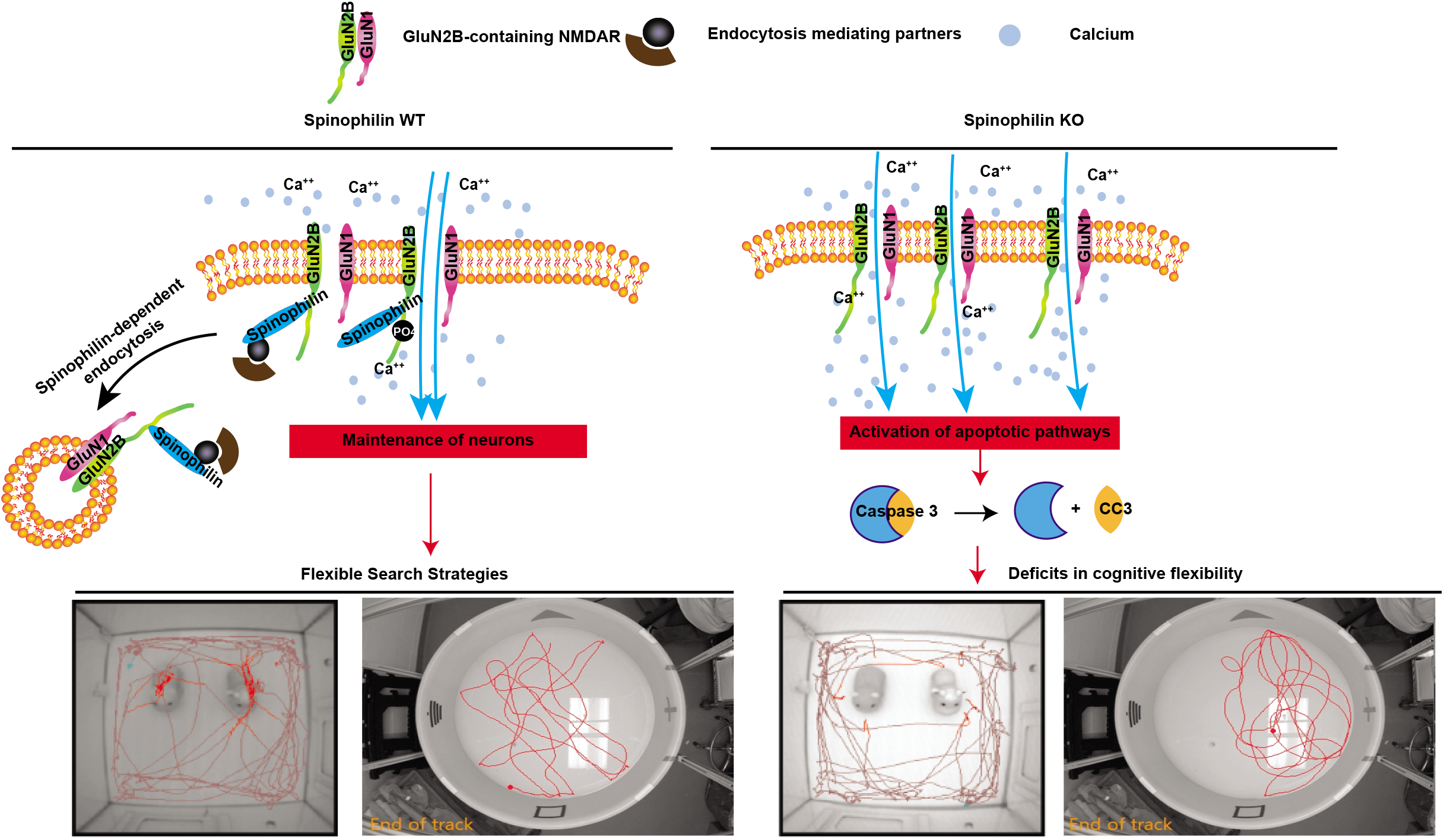

## INTRODUCTION

Postsynaptic ionotropic glutamate receptors such as N-methyl-D-aspartate receptors (NMDARs) are required for calcium influx into the postsynapse to activate signaling cascades that respond to presynaptic glutamate release. The abundance and/or calcium permeability of NMDARs at the postsynaptic membrane help to dictate the responsivity of the neuron to this presynaptic glutamate. The NMDAR is made up of 2 obligate GluN1 subunits and 2 GluN2 or GluN3 subunits. There are multiple different GluN2 subunits, with GluN2A and GluN2B having prominence in the forebrain (Paoletti et al., 2013). There are developmental and brain-region differences in isoform expression, with the GluN2B isoform being predominant at younger ages (Paoletti et al., 2013). GluN2B is also highly expressed in the hippocampus of young (~P0-P14) mice. GluN2B has a slower decay time compared to GluN2A-containing NMDARs (Paoletti et al., 2013), permitting more calcium influx into the cell. Therefore, modulation of GluN2B receptor function can robustly increase or limit calcium influx and subsequently regulate activation of signaling cascades within the postsynapse. While increases in calcium via GluN2B-containing NMDARs are critical for activation of signaling molecules such as calcium/calmodulin-dependent protein kinase II (Tavalin and Colbran, 2017), too much calcium influx can activate cleavage of apoptotic factors such as caspase-3 (Affaticati et al., 2011; Sanelli et al., 2007). Therefore, an appropriate balance of GluN2B-containing NMDAR receptor expression and activity at the postsynapse is critical for normal neuronal function and imbalances in this receptor functionality can lead to pathologies, such as activation of apoptotic pathways.

GluN2B-containing NMDAR channel activity and subcellular localization is enhanced by phosphorylation of the channel at Ser-1166 or Ser-1303 (Liao et al., 2001; Murphy et al., 2014; Tavalin and Colbran, 2017; Tu et al., 2010). Conversely, dephosphorylation of GluN2B at Ser-1480 by protein phosphatase 1 (PP1) causes translocation of the NMDAR to the synaptic fraction from the extrasynaptic fraction (Chiu et al., 2019). Therefore, multiple phosphorylation sites and mechanisms can modify GluN2B-containing NMDAR function. Spinophilin has been shown to decrease the activity of both NMDARs and AMPARs (Allen et al., 2006; Feng et al., 2000); however, the mechanisms by which spinophilin decreases NMDAR activity is unclear. Recently, we have found that PP1 can associate with the GluN2B-subunit of the NMDAR and dephosphorylate GluN2B at Ser-1284 (Salek et al., 2019). Moreover, we found that the major synaptic PP1-tageting subunit, spinophilin, limits PP1 binding to GluN2B and promotes Ser-1284 phosphorylation (Salek et al., 2019); however, functional and downstream implications of spinophilin-dependent regulation of GluN2B-containing NMDARs is lacking.

Herein, we report that Ser-1284 phosphorylation promotes GluN2B function. However, spinophilin decreases GluN2B surface expression independent of its ability to promote Ser-1284 phosphorylation. While spinophilin has multiple actions on GluN2B, loss of spinophilin enhances GluN2B surface expression in hippocampal slices, increases caspase-3 cleavage in hippocampal neuron cultures, and limits cognitive flexibility.

## RESULTS

### Spinophilin decreases NMDAR-dependent calcium influx in Neuro2a cells independent of Ser-1284 phosphorylation

The mechanisms by which loss of spinophilin stabilizes NMDAR currents by preventing current rundown (Allen et al., 2006; Feng et al., 2000) are unclear. Therefore, we tested the hypothesis that spinophilin limits calcium influx through GluN2B-containing NMDARs by transfecting NMDARs, spinophilin, and a calcium reporter, in a heterologous cell system (**Figure 1A**). To limit excessive calcium influx that leads to death upon transfection of a functional channel containing GluN1 and GluN2 subunits (Anegawa et al., 1995; Collett and Collingridge, 2004), we maintained the competitive NMDAR antagonist, AP5, in the culture media. We observed a significant decrease in GCaMP6s fluorescence in the presence, compared to the absence, of overexpressed, WT spinophilin, suggesting that spinophilin decreases calcium influx through GluN2B-containing NMDARs (**Figure 1B**). As a control, we transfected cells with only GCaMP6s in the presence or absence of spinophilin, but not NMDARs and we observed no significant increase in fluorescence upon addition of CaCl2 when the NMDAR was not transfected compared to the presence of the NMDAR. This suggests that our observation in **Figure 1B** is NMDAR dependent. To ensure that the effect is not due to autofluorescence, NMDAR subunits were expressed with and without spinophilin in the absence of GCaMP6s overexpression. The results show no significant change in the fluorescence level in these conditions suggesting that the observed result is not due to changes in autofluorescence (**Figure 1B**). As an additional control, the cells were treated with CaCl2 or vehicle. Consistent with a specific effect of extracellular calcium, our results demonstrate that the changes in the fluorescence level is solely present upon addition of extracellular CaCl2 (**Figure 1C**); however, we cannot rule out that extracellular calcium influx via NMDARs is leading to subsequent calcium release from intracellular stores which may be enhancing the GCaMP6s signal. Together, these results indicate that spinophilin significantly decreases calcium influx through GluN2B-containing NMDARs.

**Figure 1.**
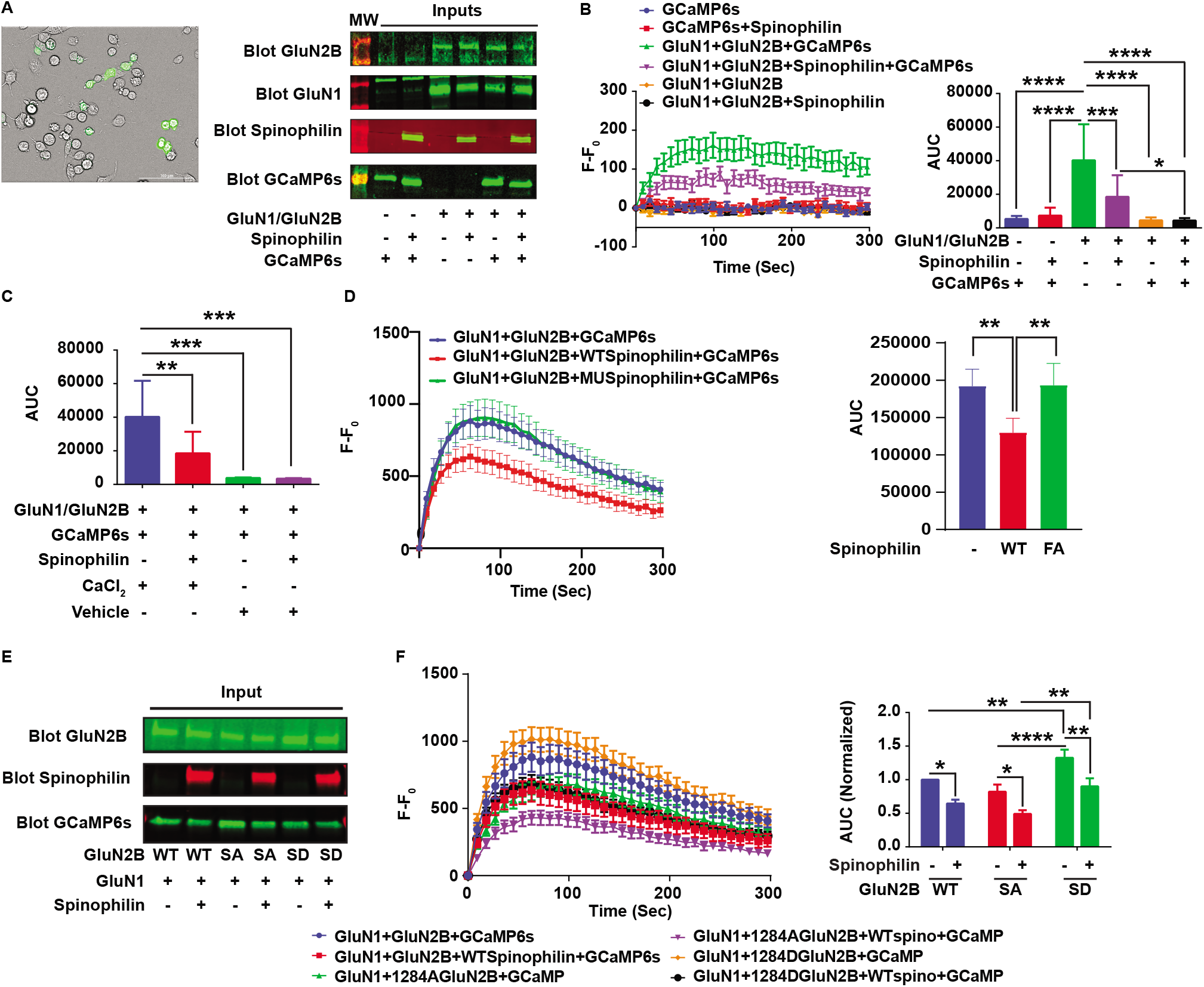
Spinophilin decreases NMDAR-dependent calcium influx in Neuro2a cells independent of Ser-1284 phosphorylation. **1A**: Brightfield and fluorescence imaging of Neuro2a cells transfected with GCaMP6s along with GluN1 and GluN2B (Left). Representative western blotting results indicating the transfection conditions and efficiency in the Neuro2a cells (Right). **1B**: Normalized (to time 0) fluorescence intensity at each time point at 9 s intervals after addition of CaCl2 to the transfected Neuro2a cells (Left) and the quantified area under the curve (AUC) indicating the total changes in the fluorescence level in each condition (Right). n=12 sets of transfections. ANOVA, F (5, 58) = 18.61, P<0.0001. **1C**: Quantified AUC of the GluN1+GluN2B with/without WT-spinophilin overexpression treated with CaCl2 or vehicle. n=12 sets of transfections. ANOVA, F (3, 30) = 10.88, P<0.0001. **1D**: Normalized (to time 0) fluorescence intensity at each time point at 9 s intervals after addition of CaCl2 to the transfected Neuro2a cells in the absence of spinophilin or the presence of WT or F451A MU spinophilin (Left) and the quantified AUC of the figure indicating the total changes in the fluorescence level in each condition (Right) n=20 sets of transfections. A matched (repeated measures) ANOVA was performed to compare samples from the same plate; F (1.372, 26.07) = 5.813; P=0.0154. Tukey Post-hoc test. **1E**: Representative western blot indicating the transfection conditions and efficiency in the Neuro2a cells transfected with different genotypes of GluN2B in the presence and absence of WT-spinophilin overexpression. **1F**: Normalized (to time 0) fluorescence intensity at each time point at 9 s intervals after addition of CaCl2 to the Neuro2a cells transfected with different genotypes of GluN2B along with GluN1 in the absence and presence of WT-spinophilin (Left) and the quantified AUC of the figure indicating the total changes in the fluorescence level in each condition (Right) n=20 sets of transfections. WT data are normalized and replotted from 1D. Two-Way ANOVA; spinophilin expression (F (1, 111) = 25.92, P<0.0001); GluN2B mutation (F (2, 111) = 13.65, P<0.0001), Interaction (F (2, 111) = 0.1553, P=0.8564). Sidak post-hoc test for spinophilin expression and Tukey post-hoc test for GluN2B mutation were performed separately. All graphs represent mean±SEM; *p<0.05, **P<0.01, ***p < 0.001 post-hoc comparisons. All the other comparisons are nonsignificant.

Next, we investigated whether spinophilin-dependent decreases in calcium influx via GluN2B-containing NMDARs is due to PP1 binding to spinophilin. To study this, we utilized a mutant spinophilin (F451A) that we and others have shown has reduced binding to PP1 (Guo et al., 2019; Hsieh-Wilson et al., 1999; Morris et al., 2018; Ragusa et al., 2010; Salek et al., 2019; Yan et al., 1999). Here, we transfected cells with GluN1, GluN2B, and GCaMP6s in the presence of wildtype (WT) or F451A mutant spinophilin. The change in GCaMP6s fluorescence upon overexpression of F451A mutant spinophilin was similar to the condition with no spinophilin overexpression (**Figure 1D**). This suggests that the spinophilin-dependent changes in calcium influx via GluN2B-containing NMDARs require PP1 binding to spinophilin.

We have previously shown that spinophilin enhances Ser-1284 phosphorylation on the GluN2B subunit of NMDARs by sequestering PP1 (Salek et al., 2019). As a result, we were interested in investigating whether spinophilin-dependent phosphorylation of Ser-1284 on GluN2B is responsible for the spinophilin-dependent decrease in calcium influx through the channel. For this purpose, we generated Ser-1284 phosphorylation-deficient (Ser to Ala) and phosphorylation-mimic (Ser to Asp) mutants of GluN2B and investigated the calcium influx through the mutant isoforms in the presence or absence of WT spinophilin. For this purpose, Neuro2a cells were transfected with GluN1, GCaMP6s, and either WT, S1284A, or S1284D GluN2B point mutants in the presence or absence of WT spinophilin (**Figure 1E**). In the absence of spinophilin, calcium influx was increased by the S1284D phosphomimetic mutation but unchanged by the S1284A non-phosphorylateable mutation. This suggests that phosphorylation of GluN2B at Ser-1284 enhances calcium influx through the channel (**Figure 1F**). However, overexpression of spinophilin in all conditions led to similar decreases in calcium influx (**Figure 1F**). Therefore, while spinophilin-dependent increases in Ser-1284 phosphorylation would be predicted to increase calcium influx through the channel, spinophilin decreases calcium influx through a pathway independent of its ability to regulate Ser-1284 phosphorylation.

### Spinophilin decreases GluN2B-containing NMDAR surface expression

To detail mechanisms by which spinophilin limits GluN2B-dependent calcium influx, surface biotinylation was used to measure surface expression of GluN1 and GluN2B subunits of the NMDAR in the presence or absence of spinophilin (**Figure 2A**). Overexpression of spinophilin significantly decreased surface expression of the GluN2B subunit (**Figure 2C**) but had no significant effect on the surface expression of the GluN1 subunit (**Figure 2B**). These experiments were performed without GCaMP6s overexpression. To ensure that changes observed in the calcium imaging experiments were not due to potential buffering or modulation of calcium binding to GCaMP6s, we repeated this study with GcAMP6s co-transfection. As above, spinophilin decreased the surface expression of GluN2B (**Figure 2E**) but not GluN1 (**Figure 2D**). As the surface levels of GluN2B were normalized to total GluN2B, these changes are not due to differences in transfection efficiency or total protein stability, but rather suggest alterations in protein trafficking. These results suggest that the spinophilin-dependent reduction in calcium influx is, at least in part, driven by enhanced internalization of GluN2B-containing NMDARs, but not GluN1 homomers, as there was no effect of spinophilin on GluN1 surface expression (see Discussion).

**Figure 2.**
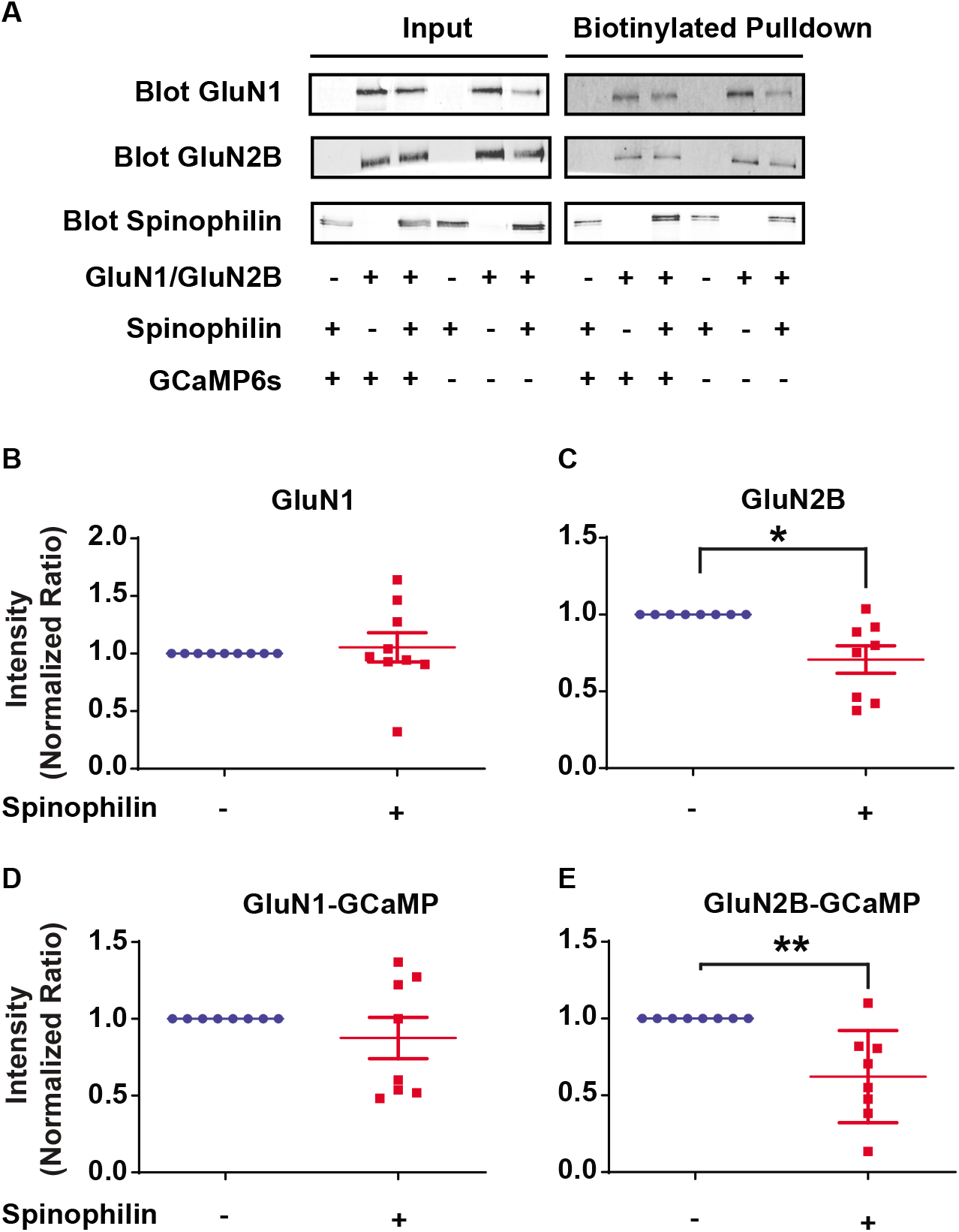
Spinophilin-dependent changes in GluN1 and GluN2B surface expression. **2A**: Representative western blots indicting the transfection conditions and biotinylation efficiency in the Neuro2a cells. **2B**: Quantified data of GluN1 surface expression in the presence or absence of WT-spinophilin overexpression with no GCaMP6s overexpression throughout the conditions. One-column t-test vs theoretical value of 1; P=0.6759. **2C**: Quantified data of GluN2B subunit surface expression in the presence or absence of WT-spinophilin overexpression with no GCaMP6s overexpression throughout the conditions. One-column t-test vs theoretical value of 1; *P=0.0135. **2D**: Quantified data of GluN1 surface expression in the presence or absence of WT-spinophilin overexpression with GCaMP6s overexpression throughout the conditions. One-column t-test vs theoretical value of 1; P=0.3847. **2E**: Quantified data of GluN2B subunit surface expression in the presence or absence of WT-spinophilin overexpression while GCaMP6s was overexpressed throughout the conditions One-column t-test vs theoretical value of 1; **P=0.0092. n=8-9 sets of transfection

### Ser-1284 phosphorylation regulates GluN2B surface expression in an activity-dependent manner

To detail if Ser-1284 phosphorylation-dependent increases in calcium-influx via GluN2B-containing NMDARs is due to increases in surface expression, Neuro2a cells were transfected with WT, S1284A, or S1284D mutant GluN2B DNA constructs along with WT GluN1 in the presence or absence of WT-spinophilin (**Figure 3A**). Overall, there was a significant decrease in surface expression in the presence, compared to absence, of spinophilin. There were no significant individual effects of spinophilin or GluN2B genotype on GluN1 surface expression (**Figure 3C**). We did not detect any significant changes in the surface expression of S1284A compared to the WT-GluN2B (**Figure 3B**). In contrast to the increased channel activity of the S128D mutant, there was a significant decrease in the surface expression of S1284D mutant (**Figure 3B**), suggesting that there is greater activity per channel rather than more channels. To test if the greater channel activity is driving an activity-dependent internalization, in addition to AP5 in the media during transfection, we performed all the biotinylation procedures with media containing AP5 to block the NMDAR activity during the biotinylation process. There was no change in the surface expression of S1284D compared to WT GluN2B when AP5 was present throughout the experiment (**Figure 3E**). Together, these results suggest that Ser-1284 phosphorylation can decrease the surface expression of GluN2B-containing NMDARs in an activity-dependent manner. Therefore, our results indicate that spinophilin driven decreases in surface expression are independent of spinophilin-dependent regulation of Ser-1284 phosphorylation.

**Figure 3.**
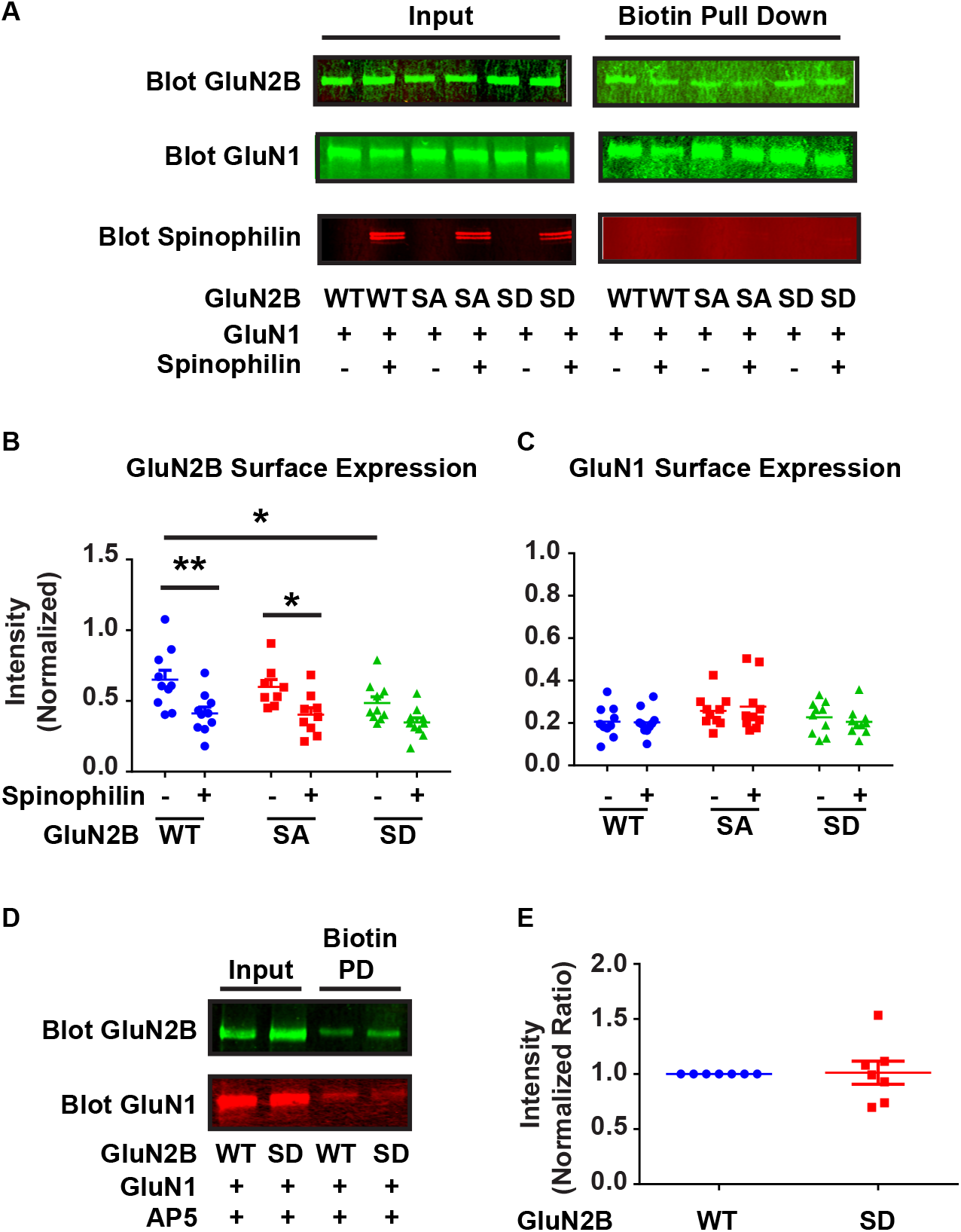
Surface expression of WT, S1284A and S1284D mutant GluN2B containing NMDA receptors. **3A**: Representative western blots indicating the transfections and biotinylation conditions and efficiency. **3B**: Quantified data indicating the surface expression of WT, S1284A, and S1284D mutant GluN2B, in the presence or absence of overexpressed WT-spinophilin. n=10 sets of transfections. Two-way ANOVA spinophilin expression (F (1, 51) = 22.36, P<0.0001); GluN2B mutation (F (2, 51) = 3.034, P=0.0569); Interaction (F (2, 51) = 0.5628, P=0.5731). Sidak post-hoc test for spinophilin expression and Tukey post-hoc test for GluN2B mutation were performed separately. **3C**: Quantified data indicating the surface expression of GluN1 when co-expressed with WT, S1284A and S1284D mutant GluN2B, in the presence and absence of overexpressed WT-spinophilin. n=10 sets of transfections. Two-way ANOVA; Spinophilin expression - F (1, 54) = 0.005796, P=0.9396), GluN2B mutation – F (2, 54) = 3.372, P=0.0417, Interaction - F (2, 54) = 0.3334, P=0.7179. No significant post-hoc differences. **3D**: Representative western blots indicating the transfection and biotinylation efficiency of WT and/or S1284D GluN2B in the presence and absence of AP5 application throughout the biotinylation procedure. n=7 sets of transfections. **3E**: Quantified data of the surface expression of WT and S1284D mutant GluN2B, in the presence of AP5 application during biotinylation n=7. One-column t-test vs theoretical value of 1; P=0.9089. All graphs represent mean±SEM; *p<0.05, **P<0.01, ***p < 0.001 post-hoc comparisons. All the other comparisons are nonsignificant.

### Surface expression of GluN2B subunit of NMDARs and GluA2 subunit of AMPARs is altered in the hippocampus of P28 spinophilin KO animals

We next investigated surface expression of NMDAR subunits in the hippocampus of WT and spinophilin KO mice. Surface biotinylation was performed on hippocampal slices generated from WT and global spinophilin KO animals. (**Figure 4A**). Consistent with overexpression studies in Neuro2A cells demonstrating that spinophilin decreased surface expression, there was a significant increase in the surface expression of the GluN2B subunit of the NMDAR (**Figure 4D**) in spinophilin KO mice. In contrast, there was no significant change in the surface expression of GluN1 (**Figure 4B**) or GluN2A (**Figure 4C**). An increased surface expression of GluN2B in spinophilin KO mice is consistent with the decreased surface expression of GluN2B upon overexpression of spinophilin in Neuro2a cells.

**Figure 4.**
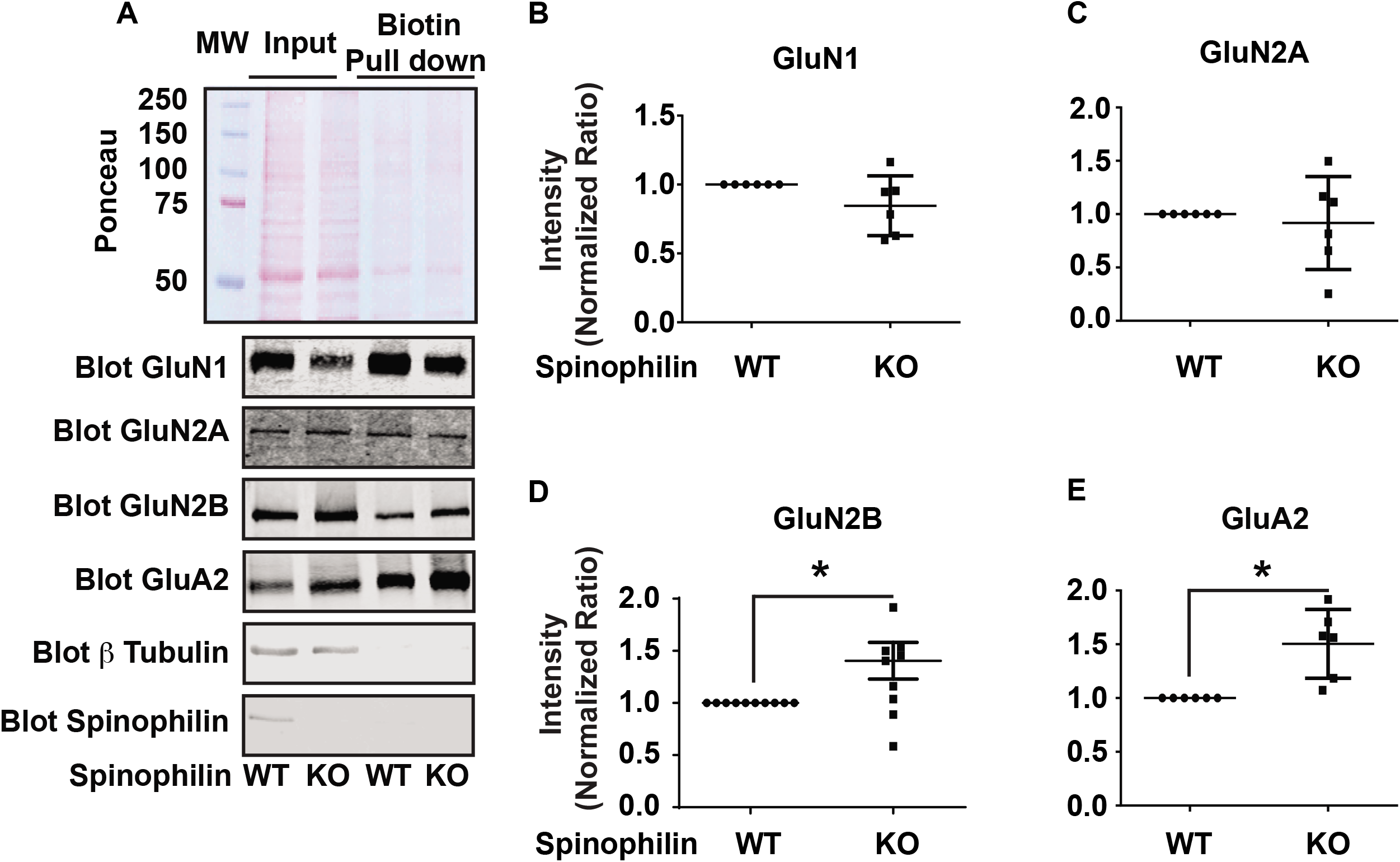
Surface expression of GluN2B subunit of NMDARs and GluA2 of AMPARs is altered in the hippocampus of spinophilin KO mouse brain. **4A**: Ponceau staining of the biotinylated inputs and pulldowns (Top). Western blotting of proteins of interest in the inputs and biotinylated pull downs. **4B**: There was no significant difference in the surface expression of GluN1 subunit. n=6. One-column t-test vs theoretical value of 1; P=0.1371. **4C**: There was no significant difference in the surface expression of GluN2A subunit. n=6. One-column t-test vs theoretical value of 1; P=0.6610. **4D**: There was a significant increase in the surface expression of GluN2B subunit. n=9. One-column t-test vs theoretical value of 1; *P=0.0371. **4E**: There was a significant increase in the surface expression of GluA2 subunit of AMPARs. n=6. One-column t-test vs theoretical value of 1; *P=0.119. All graphs represent mean±SEM.

We also investigated surface expression of the calcium-impermeable GluA2 subunit of the AMPAR (Man, 2011) given its role in regulating calcium permeability and trafficking of AMPARs. Interestingly, our results show a significant increase in the surface expression of GluA2 in the spinophilin KO hippocampi (**Figure 4E**) suggesting a potential role of spinophilin in regulating both AMPAR and NMDAR trafficking.

### The subcellular localization of NMDAR subunits is modified in P28 spinophilin global KO mouse hippocampus

To detail if loss of spinophilin impacts NMDAR subcellular localization, we used a crude fractionation protocol (**Figure S1**) to evaluate the levels of GluN1, GluN2A, and GluN2B subunit of NMDARs in S2 (membrane-associated non-postsynaptic density (PSD)) and S3 (Synaptic, PSD) fraction of postnatal day (P) 28 hippocampus of spinophilin WT and KO mice. To validate this crude fractionation, we blotted our samples for GAPDH, an S1 (cytosolic) marker, mGluR5, an S2 marker, and PSD95, an S3 marker (**Figure 5A**). Our results show a significant decrease of GluN1 in the S2 fraction but no significant change in the S3 fraction (**Figure 5B**). GluN2A results show no significant change in the S2 fraction with a significant increase in the S3 fraction (**Figure 5C**). Like GluN1, GluN2B expression was decreased in the S2 fraction but had no significant change in the S3 fraction (**Figure 5D**). These results show that global KO of spinophilin alters subcellular localization of NMDA receptor subunits in P28 mouse hippocampus. Furthermore, GluN2B-containing NMDARs are less associated with the S2 fraction, which may reflect decreases in an internalized, vesicle pool, whereas GluN2A-containing NMDARs are more associated with a synaptic fraction. Taken together with data in **Figure 4**, this suggests greater expression of GluN2B-containing NMDARs on the surface, and equal GluN2A-containing NMDARs on the surface, but a redistribution of those receptors to the synaptic fraction in spinophilin KO mice.

**Figure 5.**
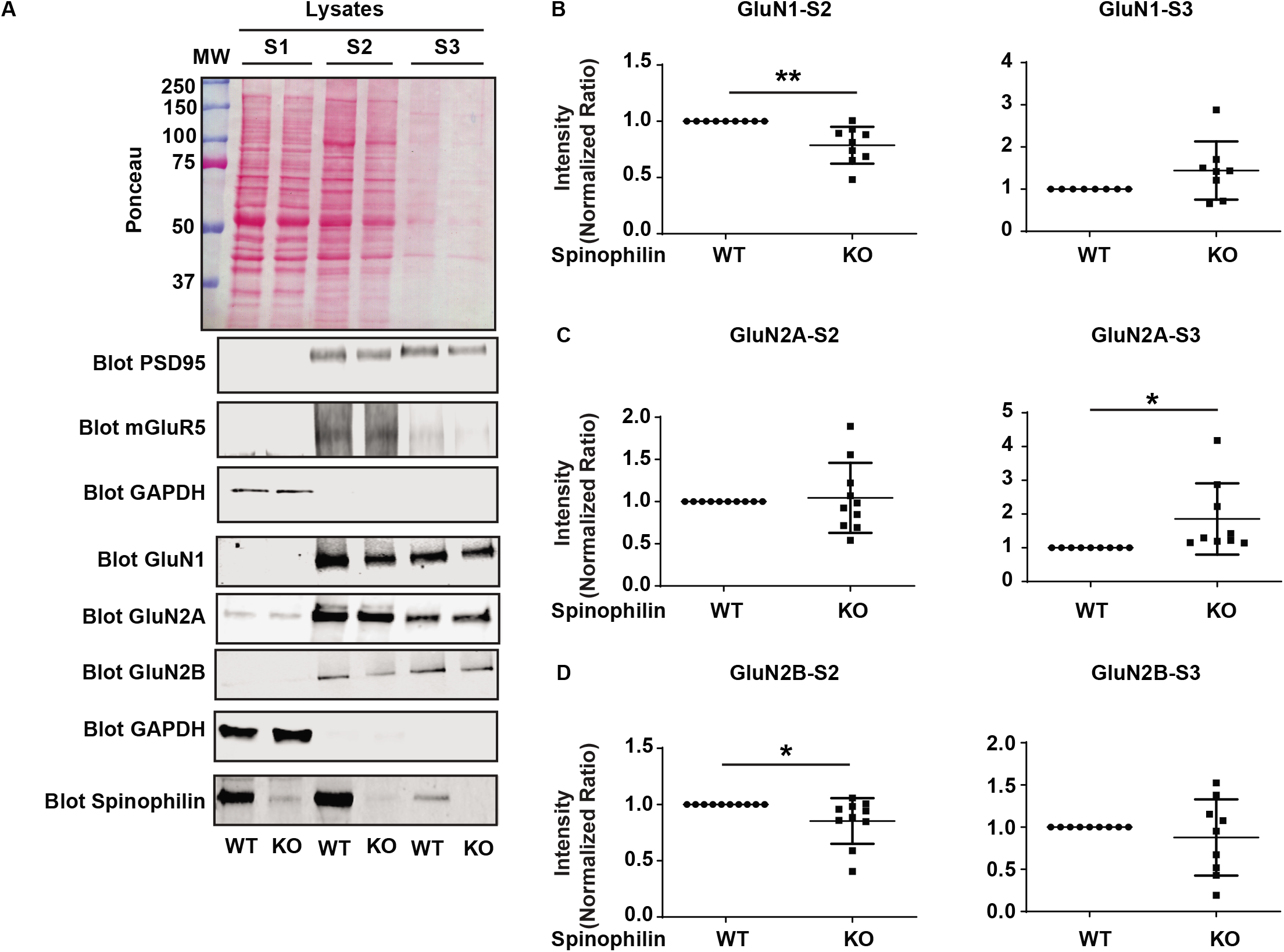
The subcellular localization of NMDAR subunits is modified in P28 spinophilin global KO mouse hippocampus. **5A**:Ponceau stain and immunoblotting of marker proteins and NMDAR subunits. **5B**: Quantified data showing the level of subcellular localization of GluN1 in S2(left) and S3 (Right) fraction. n=9. One-column t-test vs theoretical value of 1; **P=0.0044 (S2) P=0.1130 (S3). **5C**: Quantified data showing the level of subcellular localization of GluN2A in S2(left) and S3 (Right) fraction. n=9. One-column t-test vs theoretical value of 1; P=0.7438 (S2) *P=0.0413 (S3). **5D**: Quantified data showing the level of subcellular localization of GluN2B in S2(left) and S3 (Right) fraction. n=9. One-column t-test vs theoretical value of 1; *P=0.0493 (S2) P=0.4403 (S3). All graphs represent mean±SEM.

### Spinophilin KO hippocampal cultures are more susceptible to activation of apoptotic pathways

Given our previous results showing the spinophilin-dependent decreases in calcium influx via GluN2B containing NMDARs, as well as increases in surface levels of the GluN2B-containing NMDARs in spinophilin KO mice, we hypothesized that spinophilin may limit activation of calcium-dependent processes, such as caspase cleavage. To test this hypothesis, we cultured hippocampal primary neurons from global spinophilin WT and KO P0 (day of birth) mouse pups. Following 14-24 days in vitro, the conditioned neurobasal media was replaced with neurobasal media containing 100 μM glutamate for 30 minutes. Following this time, the original conditioned media was replaced, and cells were incubated for 90 minutes (**Figure S2**) and immunoblotted for cleaved caspase 3 (CC3) (**Figure 6A**). We found that the CC3 to caspase 3 (C3) ratio was three times higher in the spinophilin KO cells compared to the WT cells (**Figure 6B**). To detail if these changes were due to glutamate addition *per se*, we performed the media change in the absence of glutamate or in the presence of glutamate and AP5. Interestingly, neither NMDAR activation nor glutamate *per se* was required for the greater caspase activation observed in the spinophilin KO mice, suggesting that caspase cleavage induced by this assay is due to either the stressor induced by removal of the conditioned media and replacement with non-conditioned media or basal differences in caspase cleavage between WT and KO cells. However, spinophilin limits this caspase cleavage as there was a global significant increase in the spinophilin KO compared to WT mice (**Figures 6 C, D**). These results suggest that spinophilin KO neurons have more CC3 compared to WT neurons. While this effect does not appear to be due to acute changes in NMDAR function or calcium influx via NMDARs, we cannot conclude that it is not due to multifaceted or chronic changes in NMDAR functionality, such as redistribution of the NMDAR to different synaptic pools. Future studies will need to detail the specific mechanisms by which spinophilin regulates caspase cleavage.

**Figure 6.**
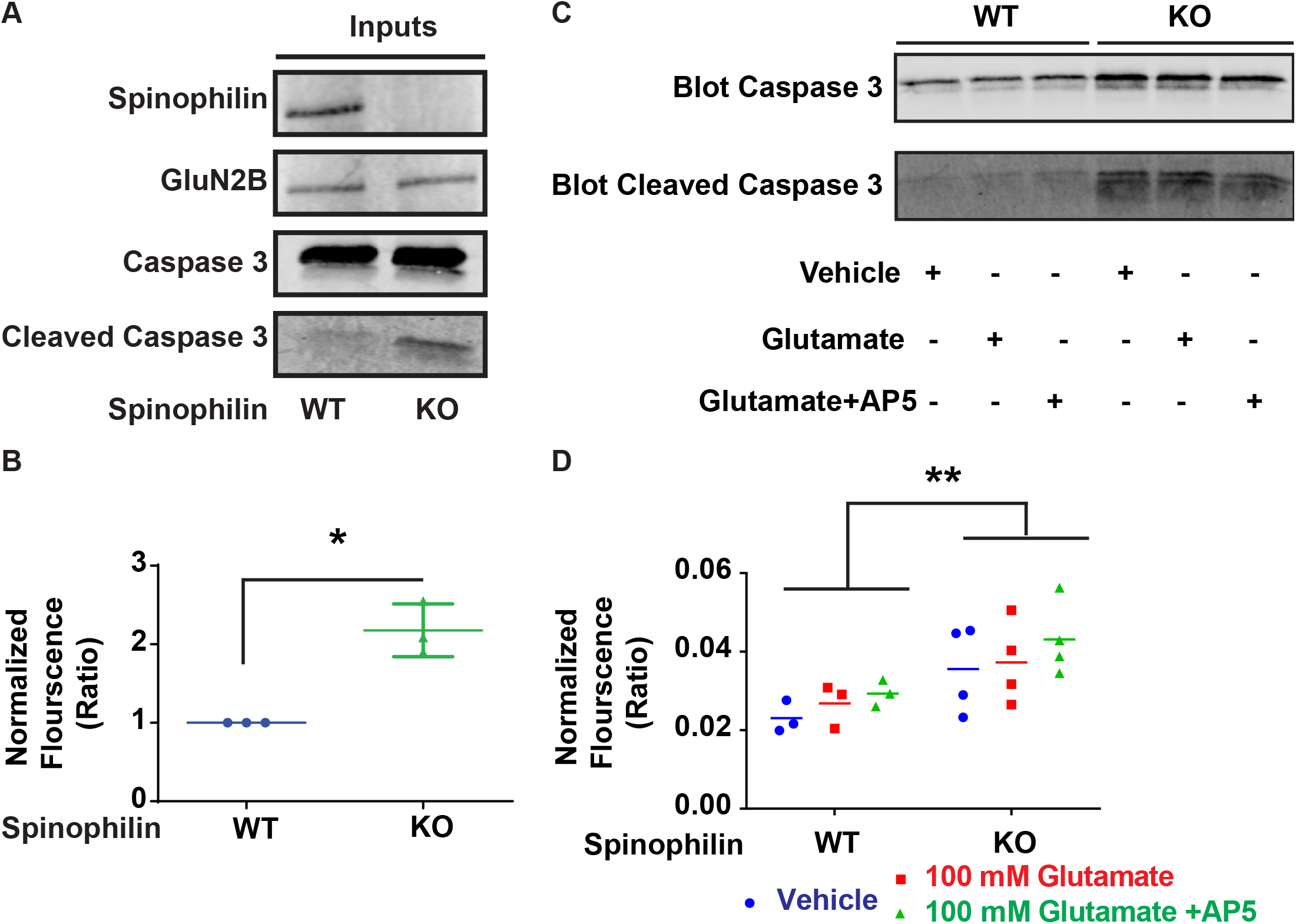
Spinophilin KO hippocampal cultures are more susceptible to activation of apoptotic pathways independent of acute calcium influx via NMDARs. **6A**: Western blot data showing C3 and CC3 bands in the spinophilin WT and KO hippocampal cultures. **6B**: Quantified data indicating a significant increase in the CC3/C3 ratio in the KOs compared to the WT cells. N=3. One-column t-test vs theoretical value of 1; *P=0.0260. **6C**: Western blot data showing C3 and CC3 bands in the spinophilin WT and KO hippocampal cultures treated with vehicle, NMDA, or NMDA and AP5. **6B**: Quantified data indicating a significant increase in the CC3/C3 ratio in the KOs compared to the WT cells. N=3. Two-way ANOVA; Spinophilin expression - F (1, 15) = 10.64, P=0.0052), Treatment – F (2, 15) = 1.147, P=0.3440, Interaction - F (2, 15) = 0.06501, P=0.9373. No post-hoc differences. All graphs represent mean±SEM.

### Spinophilin KO mice have deficits in cognitive flexibility on spatial memory tasks

While adult spinophilin KO mice may have deficits in hippocampal-dependent anxiety measures (Wu et al., 2017a) the effect of loss of spinophilin on hippocampal learning and memory are unclear. We evaluated hippocampal-dependent learning in young (starting at P28-P30) mice to limit any compensatory changes that may occur due to whole-body, constitutive loss of spinophilin. We performed a battery of 4 different behavioral tests in series (see methods), a novel location recognition (NLR) test, a novel object recognition (NOR) test, a Morris water-maze (MWM) test, and a reversal learning Morris water-maze (rMWM) test (**Figures S3 and S4**). Both WT and KO mice had similar responses to the NLR with both showing significantly more exploration of the novel location at the 30 minute, but not 24-hour time point (**Figure 7A**). In contrast, the WT mice had a trend or significantly greater exploration of the novel object in the NOR test at 30 minutes and 24 hours; however, the KO had no difference in exploration and had a trend for exploring the known object more than the novel object at 24 hours. To further test hippocampal learning, the same mice were trained on a MWM and rMWM (**Figure S4**). There were no differences in the latency of the mice to enter the quadrant where the platform was located. If anything, the KOs had a trend for a decreased latency, suggesting an increased learning (**Figure 7C**). However, when evaluating the time spent in each quadrant, the WT mice did not stay in the platform quadrant (NE) and spent the most time in the SE quadrant. In contrast the KO mice tended to perseverate in the NE quadrant (**Figure 7D**). When mice were trained on a platform in a new location (SW) for the rMWM, both mice were able to learn and had similar latencies to the new quadrant (**Figure 7E**); however, again the KO mice tended to perseverate in the initial quadrant (NE) whereas the WT mice spent equal time in all quadrants (**Figure 7F**). While there were no major differences between spinophilin WT and KO mice, there were some subtle, statistically significant, differences that suggest that the KO mice may perseverate on an initial learned task.

**Figure 7.**
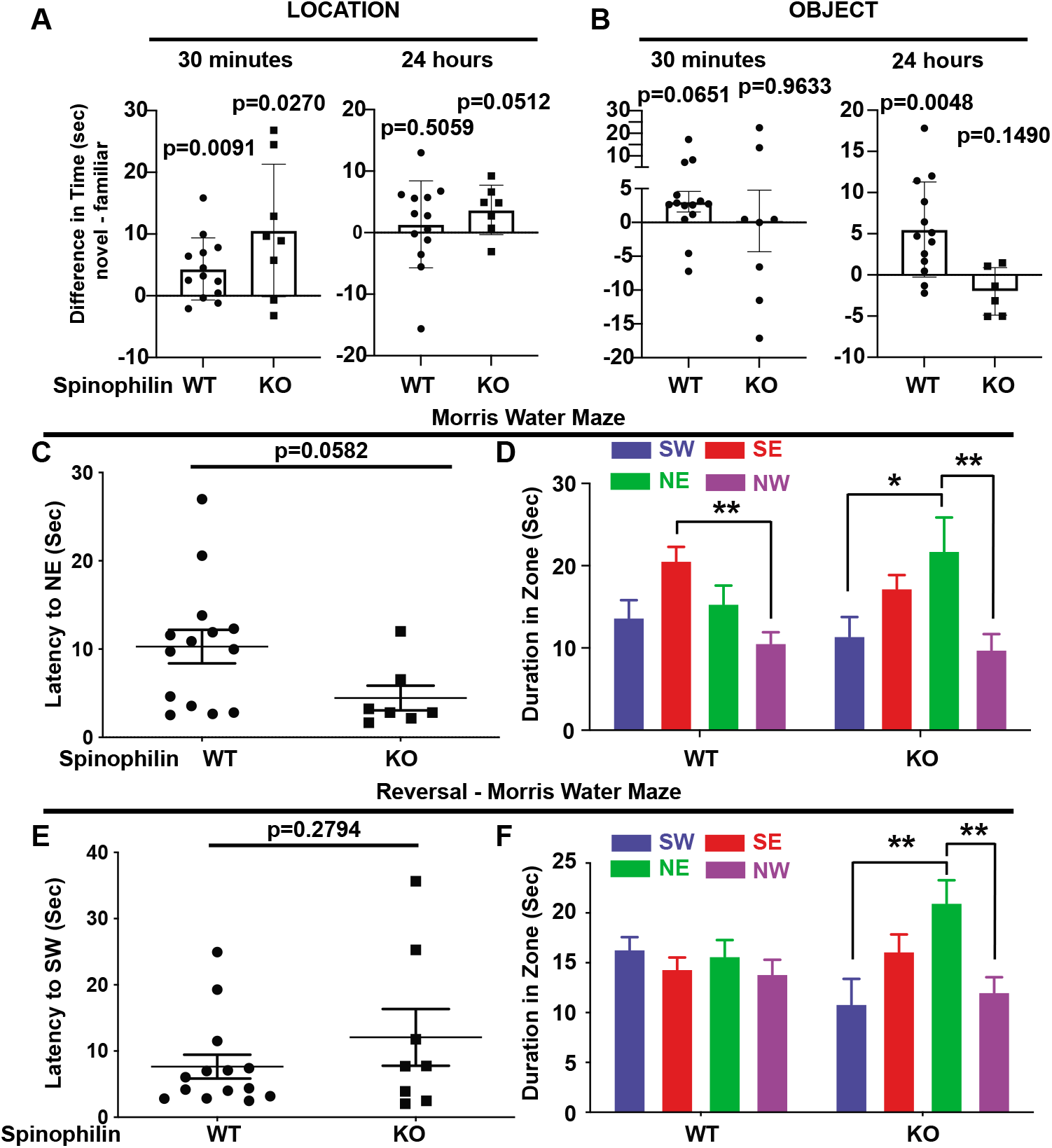
Spinophilin KO mice have normal hippocampal-dependent learning, but deficits in cognitive flexibility. **7A**: Difference in time exploring the object in the novel location, 30 minutes or 24 hours following the initial exposure to the objects. The P-value for a one-column t-test vs theoretical value of 0 is shown. **7B**: Difference in time exploring the novel compared to the familiar object, 30 minutes or 24 hours following the initial exposure to the objects. The P-value for a one-column t-test vs theoretical value of 0 is shown. **7C**: The latency to the platform quadrant (NE) between WT and KO animals. A Student’s t-test value is shown. **7D**: Time spent in the different quadrants on the probe trial day. Two-way ANOVA; Spinophilin expression - F (1, 80) = 1.012e-006, P=.9992), Quadrant - F (3, 80) = 7.153, P=0.0003, Interaction – F (3, 80) = 1.824, P=0.1495. Tukey post-hoc test shows differences between zones within the genotypes. **7E**: The latency to the platform quadrant (SW) during the rMWM test between WT and KO animals. A Student’s t-test value is shown. **7F**: Time spent in the different quadrants on the reversal probe trial day. Two-way ANOVA; Spinophilin expression - F (1, 80) = 0.0008544, P=.9768), Quadrant – F (3, 80) = 3.815, P=0.0131, Interaction - F (3, 80) = 3.590, P=0.0172. *p<0.05, **P<0.01, ***p < 0.001 Tukey post-hoc comparisons. All the other comparisons are nonsignificant.

## DISCUSSION

Spinophilin is the most abundant PP1 binding protein in the PSD (Allen et al., 1997; Colbran et al., 1997). Through this interaction, spinophilin can modulate the phosphorylation state of various proteins by either targeting PP1 activity towards or inhibiting PP1 activity at various substrates. We have found multiple subunits of the NMDA receptor in spinophilin immunoprecipitates (Baucum et al., 2013; Hiday et al., 2017; Watkins et al., 2018) and demonstrated that spinophilin interacts with the intracellular tail region of the GluN2B subunit of the NMDAR (Salek et al., 2019). Spinophilin has been shown to modulate the activity of NMDARs such that inhibition of PP1 in the presence, but not absence, of spinophilin increases NMDAR currents (Feng et al., 2000). Previous studies investigating the effect of spinophilin on NMDAR function, have not evaluated the mechanisms by which spinophilin impacts channel function. Our data indicate that spinophilin decreases calcium influx via GluN2B-containing NMDARs in Neuro2a cells. This observation can be explained in two possible ways: 1) spinophilin decreases the channel conductance to calcium and/or 2) spinophilin decreases surface expression of the receptor. The receptor surface expression was tested by quantification of surface expressed GluN2B subunits in presence and absence of spinophilin. Surface biotinylation reveals that spinophilin overexpression significantly decreases the surface expression of GluN2B, but not GluN1, in Neuro2A cells. As GluN1 and GluN2B form a functional tetramer, it may be surprising that GluN1 was unchanged. While GluN1 homomeric channels may exist at the membrane, they lack the glutamate binding site that is present in the heteromeric channel (Furukawa et al., 2005). Therefore, our data suggest that spinophilin does not regulate GluN1-containing homomers. Alternatively, as GluN1 makes up 50% of the GluN1/GluN2B heteromers, spinophilin may actually enhance homomeric GluN1 membrane trafficking and this increase may “wash out” the decrease trafficking observed in the heteromic GluN1/GluN2B channel. Future studies will need to evaluate if spinophilin can regulate GluN1/GluN2A or other functional NMDA receptor subunits or if these effects are specific to GluN2B-containing NMDA receptors. In addition to the *in vitro* effects of spinophilin, we observed increased surface expression of GluN2B in hippocampal slices isolated from global spinophilin KO animals compared to their WT control littermates. Moreover, consistent with a specific role for spinophilin on GluN2B-containing NMDAR surface expression, we observed no significant difference in GluN1 or GluN2A surface expression. Together, these data suggest that the spinophilin-dependent decrease in the GluN2B-containing NMDAR calcium influx is in part due to spinophilin-dependent decreases in surface expression. However, as stated above, we cannot rule out the possibility of spinophilin modifying channel conductance to calcium.

Spinophilin may modify GluN2B channel function independent of its ability to regulate GluN2B-phophsorylation and trafficking. We have previously shown that spinophilin promotes phosphorylation of Ser-1284 on GluN2B by decreasing PP1 association with GluN2B both *in vitro* and *in vivo* (Salek et al., 2019). We found that the S1284D phosphorylation mutant enhanced calcium influx in the GluN2B-containing NMDAR transfected cells. This increased calcium influx was not due to greater surface expression as this mutant actually had decreased surface expression compared to WT. These data suggest that our previously reported spinophilin-dependent increases in Ser-1284 phosphorylation, are not responsible for the spinophilin-dependent decreases in GluN2B surface expression, but may promote channel function. Consistent with this, overexpression of spinophilin decreases calcium influx to the same extent across the WT, S1284A, and S1284D GluN2B genotypes. Moreover, as S1284D enhances calcium influx concurrent with decreased GluN2B surface expression, these data suggest that 1284 phosphorylation enhances the calcium influx through the receptor. However, decrease in the surface expressed receptors is probably due to the greater channel function driving an activity-dependent internalization. Together, these data suggest that spinophilin can bidirectionally modulate calcium influx through NMDARs by 1) enhancing the Ser-1284 phosphorylation which increases calcium influx through GluN2B-containing NMDARs and 2) decreasing the GluN2B-NMDAR dependent calcium influx by decreasing surface expression of the channel through an unknown, activity- and Ser-1284 phosphorylation-independent pathway.

Mechanistically, how spinophilin decreases surface expression could involve clathrin-mediated endocytosis given that we have previously observed clathrin in spinophilin co-IPs (Watkins et al., 2019) and GluN2B-containing NMDA receptors undergo clathrin-mediated endocytosis (Wu et al., 2017b). Moreover, alterations in vesicle trafficking could involve the motor protein, myosin-Va, since we have previously found that both spinophilin and GluN2B interact with Myosin-Va (Baucum et al., 2013; Hiday et al., 2017; Salek et al., 2019; Watkins et al., 2018) and others have shown that myosin-Va associates with vesicles and is critical in AMPA receptor trafficking and function (Correia et al., 2008; Rudolf et al., 2011). Furthermore, in our previous study, we observed a decreased interaction of GluN2B with myosin-Va in spinophilin KO mice (Salek et al., 2019), which when taken together with decreases in GluN2B in non-PSD membranes may suggest decreased association of GluN2B with synaptic vesicle membranes in spinophilin KO mice. We previously found that other sites on GluN2B, such as Ser-929/930, Ser-1050, and Ser-1303 were not regulated by PP1 and spinophilin (Salek et al., 2019). However, additional sites on GluN2B could modulate interactions with vesicle trafficking proteins. For instance, previous studies have found that phosphorylation at Ser-1323 on GluN2B can enhance NMDA currents; however, if it modulates NMDA receptor trafficking is unclear (Liao et al., 2001). Additionally, Ser-1480 which is a PP1 site within the PDZ ligand and maintains GluN2B at extrasynaptic sites in the membrane (Chiu et al., 2019), may modulate interactions with vesicle trafficking proteins. However, if spinophilin modulates Ser-1323 or Ser-1480 phosphorylation is unknown. In addition to GluN2B, spinophilin may impact vesicle trafficking protein phosphorylation which could modulate how these proteins interact with GluN2B (and possibly other receptors that we observed similar changes in such as GluA2). Therefore, future studies need to delineate if spinophilin specifically modulates GluN2B targeting to endocytic vesicles and the mechanisms by which it does this.

Excessive calcium influx through GluN2B-containing NMDARs during glutamate toxicity plays an important role in activation of apoptotic pathways. Given that 1) excessive calcium influx can activate caspase cleavage and links to apoptosis (Affaticati et al., 2011; Sanelli et al., 2007), 2) GluN2B-containing NMDARs in general (Li et al., 2007; Liu et al., 2007; Wei et al., 1997; Werling et al., 1993), 3) Ser-1284 phosphorylation, specifically, are associated with pathological changes that lead to activation of apoptotic pathways and/or cellular toxicity (Ai et al., 2017; Li et al., 2007; Lu et al., 2015; Wei et al., 1997; Werling et al., 1993), 4) that spinophilin can decrease the calcium influx through GluN2B containing NMDARs and 5) that previous studies have linked spinophilin expression to neuroprotection and hippocampal size (Allen et al., 1997; Barui et al., 2020; Feng et al., 2000), we hypothesized that spinophilin may play a neuroprotective role. We observed greater caspase 3 cleavage in stressed hippocampal cultures isolated from spinophilin KO compared to WT mice. This effect was not due to acute activation of glutamate receptors as addition of glutamate to the stressor did not increase the activation of caspase-3, nor did AP5 block the effect. Therefore, mechanistically, it is unclear why spinophilin KO mice are more susceptible to these perturbations. However, the subcellular localization of NMDAR subunits, specifically the GluN2B subunit, plays an important role in defining the downstream signaling pathways upon activation of NMDA receptor. Specifically, activation of synaptic GluN2B-containing NMDARs is tied to pro-survival pathways while activation of extrasynaptic GluN2B NMDARs is more linked to prodeath pathways (Forder and Tymianski, 2009; Hardingham and Bading, 2002, 2010; Hardingham et al., 2002); therefore, regulation of glutamate receptor activity may induce complex changes depending on where the receptors are localized, which may activate both pro and anti-apoptotic pathways. It is unclear how spinophilin modulates NMDAR subcellular localization and if these changes directly link to the observed increases in caspase-3 cleavage. In our crude subcellular fractionation results, we found that there is a decrease in the localization of GluN2B in the extrasynaptic fraction with no significant change in the synaptic fraction. In contrast we observed increases in GluN2A in the synaptic fraction. Activation of extrasynaptic NMDARs is associated with pro-survival pathways, which is the opposite of our observations with the caspase data which show that the spinophilin KO hippocampal cultures are more susceptible to caspase 3 cleavage, a marker of apoptosis. However, as we performed a crude fractionation, this fraction also includes synaptic vesicle membranes and therefore, this decrease may be due to decreases in glutamate receptors on internalized vesicles rather than extrasynaptic GluN2B receptors. Future studies will require more sensitive high-resolution microscopy or biochemical analyses to detail how spinophilin regulates GluN2B and other NMDAR subunit localization in specific subcellular fractions. However, these data are the first to demonstrate that spinophilin KO mice have greater caspase-3 cleavage, potentially explaining why these animals have smaller hippocampi compared to WT animals (Allen et al., 1997; Feng et al., 2000).

While multiple studies have shown that loss of spinophilin impacts striatal-dependent behaviors such as locomotor and other responses to drugs of abuse, conditioned taste aversion, and rotarod coordination and learning (Allen et al., 2006; Areal et al., 2019; Edler et al., 2018; Lu et al., 2010; Morris et al., 2018; Stafstrom-Davis et al., 2001), less is known about spinophilin function in hippocampal behaviors. One recent study suggested spinophilin may be implicated in anxiety behaviors in adult mice (Wu et al., 2017a). While spinophilin is critical for regulation of cortico-striatal long-term depression (LTD), spinophilin is not required for AMPAR-dependent LTD in the hippocampus, whereas its homolog, neurabin, is (Gao et al., 2018). However, recent studies have found that mGluR5-dependent LTD in the hippocampus requires spinophilin (Di Sebastiano et al., 2016). To detail if spinophilin is required for appropriate hippocampal-dependent learning and memory, we performed a battery of 4 learning and memory tasks. While loss of spinophilin did not appear to impact the learning of the specific task, it did seem to lead to perseveration on the originally exposed object or learned task. These data may implicate spinophilin in hippocampal-dependent cognitive flexibility. For instance, hippocampal adult neurogenesis that occurs in the dentate gyrus has been found to be important in pattern discrimination and episodic memory, depression, and anxiety and attention (Gross, 2000). Moreover, these neurons are also suggested to be critical drivers of cognitive flexibility (Anacker and Hen, 2017; Weeden et al., 2019) and inhibition of NMDARs or GluN2B activity specifically has been shown to promote adult hippocampal neurogenesis and spatial memory retraining (Cameron et al., 1995; Gruden et al., 2018). Furthermore, spinophilin has been shown to interact with doublecortin (Dcx), a microtubule-associated protein, which in known to bundle microtubules in the growth cone and also is a marker of newly born neurons (Friocourt et al., 2003; Tsukada et al., 2003). Therefore, while spinophilin may have limited roles in hippocampal-dependent learning, it may be associated with cognitive flexibility that is controlled by adult-born neurons. However, motor perseveration has also been shown to be mediated by striatal function, so we cannot rule out spinophilin function in other brain regions on these behavioral changes. Future studies will need to detail specific function of spinophilin within adult-born hippocampal neurons to further test this novel hypothesis.

## SUMMARY

Together, our data demonstrate that spinophilin can bidirectionally regulate calcium influx through GluN2B-containing NMDARs, by increasing Ser-1284 phosphorylation and also decreasing GluN2B surface expression independent of Ser-1284 phosphorylation. Spinophilin limits cleavage of caspase 3 independent of acute changes in NMDAR trafficking in hippocampal neurons. Moreover, spinophilin enhances cognitive flexibility in a spatial learning and memory task, possibly suggesting a unique role for spinophilin in cells associated with these tasks, such as adult-born hippocampal neurons.

## Supporting information

All Supplemental Files

## ACKNOWLEDGEMENTS

Support for these studies comes from R33DA041876 (AJB). These studies were supported by Stark Neurosciences Research Institute, Eli Lilly and Company, and by the Indiana Clinical and Translational Sciences Institute via a Pre-doctoral fellowship to ABS, funded in part by grant # UL1TR001108 from the National Institutes of Health, National Center for Advancing Translational Sciences. The content is solely the responsibility of the authors and does not necessarily represent the official views of the National Institutes of Health. We thank Dr. Brady Atwood, Department of Pharmacology and Toxicology, Indiana University School of Medicine for critical evaluation of the manuscript.

## STAR METHODS

### Reagents

All custom materials will be shared upon reasonable request. Experiments were approved by the institutional biosafety committee (IBC-1594 and IN-1000).

cDNAs: Expression vectors used in this study including human WT and F451A mutant spinophilin and C-terminal tail of GluN2B, were previously described (Hiday et al., 2017; Salek et al., 2019). Human full-length GluN2B (BC113618; Transomic Technologies, Huntsville, AL, USA) or mouse GluN1 cDNA (BC039157; Transomic Techologies) were shuttled into an expression vector with an N-terminal myc or V5 tag, respectively. GCaMP6s expression vector was obtained from Addgene (#40753, Watertown, MA, USA). Transfection Reagent: Polyjet (SignaGen Laboratories, Rockville, MD, USA) was used for transfections.

Antibodies: Antibodies used for IPs and/or primary blotting:

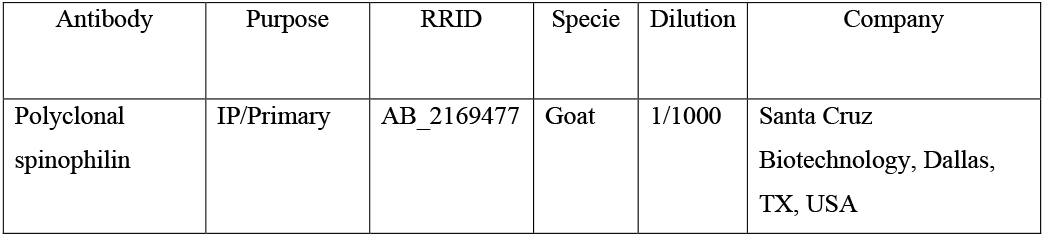
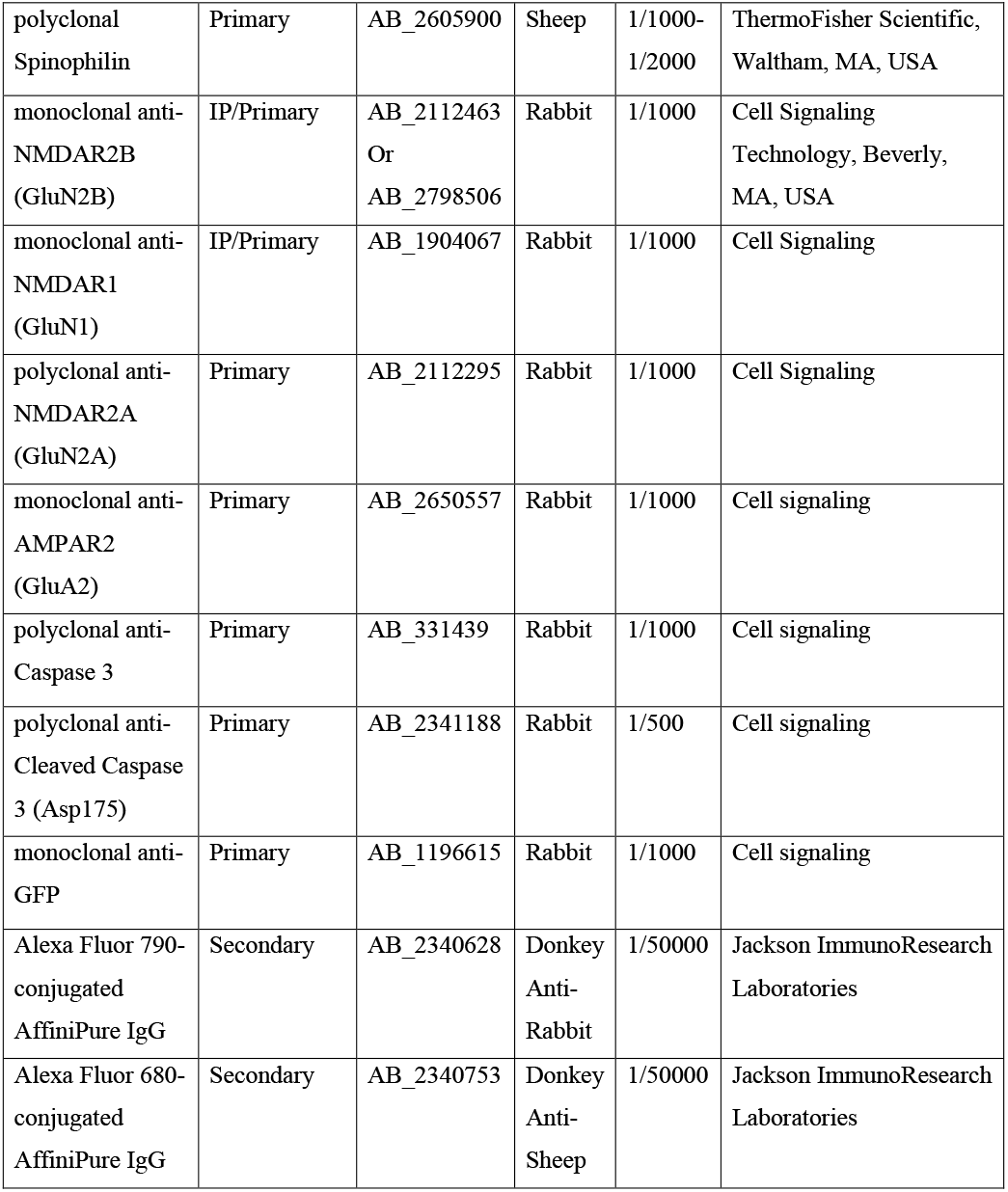

Other reagents: D-AP5 (D-145, Alomone, Jerusalem, Israel) or (14539, Cayman Chemicals, Ann Arbor, MI, USA), BSA (A9647-100G, Sigma-Aldrich, St Louis, MO, USA), Leibovitz’s L-15 media (21083027, Gibco by Life Technologies, ThermoFisher Scientific, Waltham, MA, USA), Papain (P4762-500MG, Sigma-Aldrich), Modified Eagle’s Medium (MEM) (51200038, Gibco by Life Technologies, ThermoFisher Scientific), Fetal Bovine Serum (FBS) (16140063, Gibco by Life Technologies, ThermoFisher Scientific), Horse serum (26050070, Gibco by Life Technologies, ThermoFisher Scientific), L-Glutamine (35050-061, Gibco by Life Technologies, ThermoFisher Scientific), Penicillin/Streptomycin (Pen/Strep) (15140-122, Gibco by Life Technologies, ThermoFisher Scientific), Glucose (G5767-500G, Sigma Aldrich), Insulin/Selenite/Transferrin (IST) (I1884-1VL, Sigma-Aldrich), Neurobasal Media (12349015, Gibco by Life Technologies, ThermoFisher Scientific), B27 Supplement (17504044, Gibco by Life Technologies, ThermoFisher Scientific), Gentamycin reagent (15750-060, Gibco by Life Technologies, ThermoFisher Scientific). Sulfo-NHS-SS-Biotin (325143-98-4, A8005, APExBIO Technology LLC, Houston, Texas, USA), Neutravidin beads (29201, ThermoFisher Scientific), Protease inhibitor cocktail (Thermo-Fisher Scientific or Bimake, Houston, TX, USA), Glutamate (G2834-100G, L-Glutamic acid, monosodium, Salt monohydrate, 98%, Molecular weight:187.13 (GA)), 12 mm coverslips (1254582, ThermoFisher Scientific), Cytosine β-D-arabinofuranoside (Ara-C) (C1768-100MG, Sigma-Aldrich), White nontoxic tempura washable paint powder (Crayola).

All other utilized reagents were of highest purity obtained from ThermoFisher Scientific, Sigma-Aldrich, or Gibco.

### Equipment

Phenotyper cages (Noldus Information Technology, Wageningen, Netherlands), Ethovision XT video tracking software (Noldus Information Technology), Toys (Spark Create Imagine), 120 cm MWM pool (Maze engineers, Cambridge, MA), Heater (Maze engineers), Platform (30cm, Maze engineers), Gantry (Maze engineers), Color camera (Noldus Information Technology).

### Animals

Experiments were approved by the School of Science Institutional Animal Care and Use Committee (SC270R, SC310R) and performed in accordance with the Guide for the Care and Use of Laboratory Animals and under the oversight of the Indiana University-Purdue University, Indianapolis (IUPUI). Both male and female mice were used and as they were below the age of sexual maturity, we pooled data together. Animals were provided food and water ad libitum. Mice were maintained on a normal 12 hour light (7 am - 7 pm) / dark (7 pm - 7 am) cycle. Spinophilin KO mice were initially purchased from Jackson Laboratories (Bar Harbor, ME, USA; Stock #018609; RRID: MMRRC_049172-UCD) and a breeding colony has been maintained at IUPUI. Male or female, WT, C57Bl6, (Jackson laboratories) or spinophilin knockout mouse brains were dissected at Postnatal day 28–32 (P28). Animals were group housed and WT and KO littermates were used (WT and KO animals were from heterozygote x heterozygote breeding pairs). Animals were weaned ~ P21. For behavior tests, Animals were group housed until the genotyping was completed ~ P23. At ~P23, male or female, spinophilin KO mice or WT littermates were singly housed throughout the behavior studies. All behavioral tests were performed between 11AM to 6PM. For generation of neuronal cultures, mice were weaned at P0. For biochemical analyses and generation of neuronal cultures, animals were euthanized by decapitation without anesthesia.

### Mutagenesis

Mutagenesis was performed as described before (Salek et al., 2019). Briefly, reactions were performed using the site-directed mutagenesis kit (Agilent Technologies, Santa Clara, CA, USA) using Q5 DNA polymerase in Q5 DNA buffer in the presence of DNTPs and template DNA followed by a mutagenesis PCR protocol. The PCR products were later transformed into DH5α E. coli. Vectors were then sequence verified (Genewiz Inc, South Plainfield, NJ, USA) for the mutations.

### Mammalian protein expression

Neuro2a cells were used for mammalian protein expression. Cells were purchased and split into passage 9 and frozen down. The cells were used up to passage 22. After thawing, cells were incubated with MEM recovery media containing 20% FBS, 1% Pen/strep and 1% Sodium Pyruvate. The cell culture and incubation after recovery was performed in MEM containing: 10% FBS, 1% Pen/Strep and 1% Sodium Pyruvate. 6- and 12-well plates were placed in a tissue culture incubator (Panasonic Healthcare; Secaucus, NJ, USA) at 37°C and 5% CO2. Cells were counted and the density was adjusted to 70,000 - 100,000 cells/mL. 6-well and 12-well plates received 2 mL or 1 mL, respectively, of cell containing media. Cells were transfected the next day at about 50-60% confluency. Confluency was measured by estimating cell coverage on the bottom of the flask. For 6-well plates, DNA (0.5–2 μg per DNA vector) was added to 250 μL of serum-free MEM in a 1.7 mL microcentrifuge tube. In a separate microfuge tube, transfection reagent was added to 250 μL of serum-free MEM. Polyjet was used in a 3: 1 volume: mass ratio (e.g. 9 μL of Polyjet was used with 3 μg DNA). For each well, DNA concentrations were equalized using an empty DNA vector, so that each condition in the same experiment had an equal mass of DNA and transfection reagent (all the volumes were cut in half for 12-well plate studies). The Polyjet containing MEM was then added to the tube containing DNA and incubated at room temperature for 15 min. The DNA-Polyjet mixture was then added to each well very slowly as the plate was being gently shaken on a horizontal axis to mix the DNA mixture with the media. The wells transfected with full-length GluN1 and full-length GluN2B expression vectors were then treated with 0.25 μg/mL of AP5. 3 μg of AP5 was dissolved in 600 μL of MEM. 100 μL of this mixture was then added to each well of a 6-well plate containing 2 mL of cell media. The cells were then placed in the incubator overnight and were processed the next day. If the cells were cultured for imaging, they were imaged the next day prior to lysis. If not, the cells were processed the next day as follows: MEM was removed, and cells were washed with 2 mL of cold phosphate-buffered saline (PBS). PBS was aspirated off and cells were lysed in 0.75 mL KCl lysis buffer (150 mM KCl, 1 mM dithiothreitol (DTT), 2 mM EDTA, 50 mM Tris-HCl pH 7.5, 1% (v/v) Triton X-100, 20 mM β-glycerophosphate, 20 mM sodium fluoride, 10 mM sodium pyrophosphate, 20 mM sodium orthovanadate, 1X protease inhibitor cocktail) then transferred into 2 mL microcentrifuge tubes. Cells were sonicated at 25% amplitude for 15 s at 4°C using a probe sonicator (Thermo-Fisher Scientific) and centrifuged (4°C for 10 min at 16,900 x g).

### Calcium imaging in Neuro2a cells

To image the changes in intracellular calcium levels, Neuro2a cells were plated in 12-well plates and were transfected with 0.5 μg each of V5-GluN1, Myc-GluN2B, GCaMP6s and/or 1 μg WT or F451A mutant HA-spinophilin. Each well received a final concentration of 2.5 μg/mL AP5 after the transfection. The next day, AP5 containing MEM was aspirated off and replaced with 1 mL of calcium-free 1X PBS in room temperature. The cells were immediately placed in the Cytation 3 (Biotek, Winooski, Vermont, U.S.A.) cell imaging multi-mode reader. The reader was temperature and gas controlled and was set at 35-37°C and 5% CO2. Changes in fluorescence were measured for 5 minutes at 9 s intervals, resulting in a total of 34 readings. To measure fluorescence, the excitation and emission wavelengths were set at 488 and 528 nm, respectively, reading from the bottom of the plate. This 5-minute incubation with calcium-free PBS, was used to minimize background fluorescence by decreasing intracellular calcium and concomitant GCaMP6s fluorescence. After the incubation, each well received CaCl2 (3-6 mM final concentration) via a built-in dispenser and briefly received an orbital shake for 2 s to uniformly mix the CaCl2 with the media. The plate was read at the same wavelengths mentioned above for another 5 minutes. After the reading was completed, the media was aspirated off the cells. The cells were then lysed and processed as above in 350 μL RIPA/PI lysis buffer (1 mM EDTA, 150 mM NaCl, 20 mM Tris-HCl, 1X protease inhibitor, 20 mM β-glycerophosphate, 20 mM sodium fluoride, 10 mM sodium pyrophosphate, 20 mM sodium orthovanadate, 0.01% NP-40, 0.01% deoxycholate). All the fluorescence reading values at each time point were normalized back to the baseline by subtracting each value from the fluorescence value at the 0-time point of the corresponding well. The data was then used to plot a graph and the area under the curve (AUC) was quantified.

### Biotinylation in Neuro2a cells

Neuro2a cell biotinylation was based on previously published protocol (Cao et al., 2007) and optimized for a 6-well cell culture plate. Neuro2a cells were washed 3 times in 1 mL of B buffer (0.5 mM CaCl2, 0.5 mM MgCl2-6H2O in 1X PBS). Following the wash, 1 mL of 0.5 mg/mL Sulfo-NHS-SS-Biotin in B buffer was added to each well and allowed to incubate at room temperature for 5 minutes. The free biotin was then quenched by washing the cells twice with 1 mL of biotin quenching buffer (100 mM Glycine in B buffer). Following the quenching step, the cells were lysed in 750 μL of RIPA/PI buffer. The samples were then centrifuged at 14,000 x g at 4°C for 15 minutes. To create the input, 150 μL of the lysate supernatant was mixed with 50 μL of 4X SDS containing sample buffer with DTT. 500 μL of the supernatant was mixed with 40 μL of a 50% slurry of NeutrAvidin beads and incubated with rotating at 4°C overnight. The next day, the beads were washed 3X by centrifuging the samples at 2000 x g for 1 minute and replacing the supernatant with 500 μL RIPA/PI buffer and allowing to rock for 5 minutes at 4°C. Following the last wash, 60 μL of 2X SDS containing sample buffer with DTT was added to the beads. The beads were thoroughly vortexed and placed at −20°C for western blotting.

### Biotinylation in brain slices

The protocol used for brain slice biotinylation is modified from a previous study (Gabriel et al., 2014). Room temperature, 1X artificial cerebrospinal fluid (aCSF; 125 mM NaCl, 2.5 mM KCl, 1.2 mM NaH2PO4, 1.2 mM MgCl2-6H2O, 2.4 mM CaCl2, 26 mM NaHCO3, 11 mM Glucose) and ice-cold high sucrose solution (HSS) (250 mM Sucrose, 2.5 mM KCl, 1.2 mM NaH2PO4, 2.4 mM CaCl2, 26 mM NaHCO3, 11mM Glucose) were prepared and bubbled with carbogen (95% O2 and 5% CO2) for a minimum of 20 minutes. Animals were decapitated and the brains were dissected on ice. The brains were quickly transferred to ice-cold HSS and 300 μM slices were generated using a VT1200-S vibrating microtome (Leica Biosystems, Buffalo Grove, IL). Four hippocampi containing whole brain slices were generated from each brain. The slices were then transferred into a slice chamber and were incubated with 31°C, circulating, carbogenated 1x aCSF for 40 minutes to recover. The following procedures were performed on ice unless otherwise stated. After the 40-minute recovery, the slices were transferred into an ice-cold, 24-well plate and were incubated with 750 μL of 1 mg/mL of Sulfo-NHS-SS-Biotin dissolved in ice-cold carbogenated 1X aCSF for 45 minutes followed by a 3X wash with 750 μL of ice-cold 1X aCSF. After the last wash, the slices were incubated with 750 μL of ice-cold 1X aCSF for 10 minutes followed by 3 washes with 750 μL of ice-cold biotin quenching buffer (100 mM Glycine in 1X aCSF). Following the last wash, the slices were incubated with 750 μL of ice-cold biotin quenching buffer for 25 mins. The slices were then washed 3X with ice-cold 1X aCSF. After the last wash, the hippocampi were dissected from the slices and were transferred into a homogenizer containing 1200 μL of RIPA/PI buffer. The slices were homogenized using 18-20 up-and-down movements of a pestle in a 2-mL tight-fitting glass homogenizer. The homogenate was then sonicated once for 15 seconds at 25% amplitude followed by centrifugation for 15 minutes at 14,000 x g at 4°C. 150 μL of the lysate supernatant was mixed with 50 μL of 4X SDS sample buffer and used as the input. 500 μL of the supernatant was mixed with 60 μL of pre-washed NeutrAvidin beads and rotated at 4°C overnight to pulldown (PD) biotinylated proteins. The next day, the PD samples were washed three times using 500 μL of ice-cold RIPA/PI buffer. The samples were centrifuged for 1 minute at 2,000 x g, then the supernatant was aspirated off and replaced with 500 μL of icecold RIPA/PI. The tubes were then replaced on the rotator and were allowed to rotate at 4°C for 5 minutes. This wash procedure was repeated 3 times. Following the last wash, the supernatant was removed and 60 μL of 2X SDS sample buffer+DTT was added to the beads, the tubes were briefly vortexed and placed at −20°C and saved for SDS-PAGE. The inputs and biotinylated PDs were separated by SDS-PAGE and immunoblotted. The intensity of the PD band of the protein of interest was normalized to the input band of the same protein, showing the level of protein surface expression. Finally, the ratio for each protein of interest in the KO samples was additionally normalized to the WT sample on the same gel.

### Subcellular fractionation

Male and female P28-P32 Spinophilin WT and KO mice were decapitated without anesthesia, brains were removed, the hippocampi were rapidly dissected, frozen in liquid nitrogen, and stored at −80 °C. Two whole frozen mouse hippocampi (one from each hemisphere) were pooled and homogenized in 2 ml of a detergent-free isotonic (150 mM KCl, 50 mM Tris-HCl, 1 mM DTT, 2 mM EDTA, 1X protease cocktail, 20 mM NaF, 20 mM NaVO4, 20 mM Beta-glycerophosphate, 20mM Na-Pyrophosphate) buffer homogenized using 18-20 up-and-down movements of a pestle in a 2 mL tight-fitting glass homogenizer. Samples were then centrifuged at 100,000 × g at 4°C for 1 hour. Supernatants (S1) were saved for SDS-PAGE. The pellet (P1) was resuspended in 1 ml of isotonic buffer containing 0.5% Triton X-100 in a microcentrifuge tube. Samples were then centrifuged at 14,000 × g at 4°C for 10 min. Supernatants (S2) were saved, and the P2 pellets were resuspended in 1 ml of isotonic buffer containing 1% Triton X-100 and 1% sodium deoxycholate and sonicated. Following incubation at 4°C for 15 seconds. samples were then centrifuged at 14,000 × g for 10 min, and the supernatants (S3) were saved for SDS-PAGE. The 150 μL of lysate samples from S1, S2, and S3 fractions were mixed with 4X SDS sample buffer containing DTT and the same amount of each sample from WT and KO brains was loaded on the gel. After the transfer of the proteins to nitrocellulose membrane, the membranes were stained with 1X Ponceau S stain for 5 minutes to stain for total proteins. After western blotting, the intensity of the band of the protein of interest was divided by the total amount of protein in the sample measured by Ponceau S stain using ImageJ. This value in the KOs was divided by that of the WT in the same trial.

### Hippocampal primary neuronal cultures

Hippocampal primary neurons were dissociated and cultured using a previously published protocol (Bansal et al., 2019). In short, hippocampi were dissected from P0 mice in harvest media (0.02% BSA in Leibovitz’s L-15 media). The hippocampi were then transferred to a 15 mL conical containing 0.5 mL dissociation media (0.038% papain in previously made 0.02% BSA/L15), carbogenated, recovered in a 37°C water bath for 10 minutes, incubated in M5-5 media (5% FBS, 5% Horse serum, 0.2% L-Glutamine, 1% Pen/Strep, 1% Glucose, 0.25% IST in MEM), and dissociated by pipetting 30X using three different sizes of sterile, fire-polished, pasture pipets. After each round of pipetting the supernatant containing the dissociated cells was collected and pooled in a sterile 15 mL conical centrifuge tube. After the last round, the supernatant was centrifuged at 800 x g for 5 minutes at room temperature. The supernatant was removed, and the pellet was dissolved in 3 mL of M5-5 media. Then, 1 mL of the cell mixture was added to each well of a 24-well plate containing a 12 mm coverslip previously coated with 0.5 mg/mL Poly-Lysine and placed in 5% CO2, 37°C cell culture incubator. After 48 hours, 0.5 mL of the M5-5 media was replaced with 1 mL of B27 supplement media (2% B27 Supplement, 0.25% L-Glutamine, 0.25% IST, 0.1% Gentamycin reagent and 15μL Ara-C in Neurobasal media, mixed and sterile filtered using 0.2 μm filter syringe). The plate was then placed back in the incubator until the day of the experiment (14-24 days in vitro; DIV).

### Hippocampal neuron stress paradigm

Primary hippocampal cultures were assayed at 14-24 DIV. On the test day, the culture media was collected from the wells into a sterile 15 ml conical centrifuge tube to generate the conditioned Neurobasal (cNB) media. Each of the spinophilin WT and KO wells received 1 mL of fresh neurobasal media alone, media with 100 μM glutamate, or media 100 μM glutamate + 2.5 μg/ml AP5. The cells were incubated with the media for 30 minutes in 37°C cell incubator. During incubation, the cNB media was sterile filtered using 0.2 μm filters and the volume was adjusted by adding fresh Neurobasal media (no more than 10% of the total media volume) and placed in a 37°C water bath. After 30 minutes, the glutamate containing media was removed from the wells and replaced with 1.5 mL cNB media and incubated in a cell culture incubator for 90 minutes to recover. Following the recovery step, the neurons were lysed and processed as follows: the media was removed from the wells and 300 μL of ice cold low ionic lysis buffer (2 mM Tris pH 7.5, 2 mM EDTA, 1 mM DTT, 1x Protease Inhibitor, 20 mM NaF, 20 mM Na orthovanadate, 10 mM Na pyrophosphate, 20 mM β-glycerophosphate, 1% Triton) was added to the cells. The cells were lysed by trituration until all the cells were detached from bottom of the wells. The lysate was then transferred into 1.7 mL microcentrifuge tubes and sonicated for 15 seconds at 25% amplitude with a probe sonicator followed by centrifugation at 4°C for 15 minutes at 14,000 x g. 150 μL of the lysate supernatant was transferred into a new tube and mixed with 4X SDS sample buffer containing DTT. The samples were then placed at −20°C until processed.

### Immunoprecipitations and Western Blotting

Cell lysate inputs and/or PD samples were used for western blotting. All samples were heated at 70°C for 10 min prior to loading on the gel. PD samples were briefly vortexed and centrifuged at 1000 x g for 2 minutes to precipitate the NeutroAvidin agarose beads and separate them from the suspension prior to loading on the gel. 10-35 μL of input or 20-30 μL of PD sample were loaded onto a 1.00 mm hand-cast 10% polyacrylamide gel. The gels were electrophoresed at 75 V for 15 min and 175 V for approximately 1 h. Proteins were transferred to a nitrocellulose membrane using a wet transfer in a CAPS transfer buffer (10% MeOH, 0.01 M N-cyclohexyl-3-aminopropanesulfonic acid pH 11). The transfer was performed in a transfer tank attached to a cooling unit set at 4°C and transfer was operated at a constant 1.0 Amps for 1.5 h. Membranes were stained with a 2 mg/mL Ponceau S stain dissolved in 10% Trichloroacetic acid for 5 min to normalize inputs for equal loading where applicable. Following Ponceau staining, membranes were scanned and subsequently washed with deionized water. Membranes were blocked in Trisbuffered saline with Tween (TBST; 50 mM Tris pH 7.5, 150 mM NaCl, 0.1% (v/v) Tween-20) containing 5% (w/v) nonfat dry milk. Blocking was performed three times, 10 min each, for a total of 30 min. Membrane was incubated with primary antibodies diluted in 5% milk in TBST overnight at 4°C with gentle shaking. After incubation, membranes were washed 3X for 10 min per wash with TBST containing 5% milk. Appropriate secondary antibodies in TBST containing 5% milk were added to the membranes following the washes. Jackson ImmunoResearch antibodies were typically diluted 1 : 50,000 and Invitrogen antibodies were generally diluted 1 : 10000. Secondary antibodies were incubated with membranes for 1 h at 22°C shaking in darkness. Membranes were washed three times with TBS without Tween for 10 min for each wash. Fluorescence scans were performed using the Odyssey imaging system (LiCor, Lincoln, NE, USA) and data analysis was done using Image Studio software (LiCor). We have previously shown linearity of fluorescence intensity using these conditions for multiple proteins and antibody pairs (Edler et al., 2018; Morris et al., 2018).

### Novel Object Recognition and Novel Location Recognition

#### Object And Location Setting

The objects were odorless plastic baby toys. The toys had no sharp edges. The toys had different shapes and shades of color. Since the toys were lightweight, they were filled with cement and allowed to completely dry before use so they cannot be moved by the mice (**Figure S3A**).

Testing arena is a square space (Phenotyper cages, dimensions: 30 x 30 cm) in which all the animal behavior is monitored by a built-in camera in the cage lid. The arena consists of the floor of the phenotyper cage (**Figure S3B**).

Testing zones are the rectangular space designated around each object. The locations of the zones are designed to have the same distance from the corners and the walls (**Figure S3C**). During the testing session, the software records the cumulative duration of time that the animal’s nose point is in each zone or outside of the zones. After the test, the number of entrances (frequency) and the cumulative duration of the times that animal’s nose point was in the zones are quantified using the software and are used as an index to study time and frequency of exploration of the novel and familiar object or location. After each use of the Phenotyper cage, the cage is sprayed and wiped down with 70% ethanol to eliminate any potential odor or residuals from the previous animal.

#### Habituation

The animals were habituated to the environment for three consecutive days, 15 minutes per day. During the habituation session, each animal was placed in the clean and empty Phenotyper cage and allowed to explore for 15 minutes (**Figure S3B**). After 15 minutes, the animal was gently removed from the cage and replaced in the home cage. The cage was sprayed and wiped down with 70% ethanol.

#### Familiarization

Two identical objects (parrots, hippos, or giraffes) were placed in the designated zones. The objects had the same distance from the walls and corners. The animals were then placed into the arena and were allowed to familiarize with the objects for 20 minutes (**Figure S3D**).

#### Acquisition

After acquisition, the animals were returned to their home cage. The testing was performed at two different time points: 30 mins post acquisition to explore short-term memory and 24 hours post acquisition to explore long-term memory.

#### Testing

Following the habituation and familiarization phase, two different testing paradigms were used: NLR and NOR

#### Novel Location Recognition-30 minutes

30 minutes post familiarization, the NLR test was performed. At the time of testing, the location of one of the familiar objects was changed. The animal was then placed in the Phenotyper cage and was allowed to explore for 5 minutes (**Figure S3E**). After the 5 minutes, the animal was returned to the home cage and allowed to recover for 5 minutes. After the test, the cumulative duration of the times in which the nose point was in the boundaries of familiar object and novel object zones and the frequency of entries to them was measured using Ethovision software.

#### Novel Object Recognition-30 minutes

While the animal was recovering in the home cage after the NLR test, the Phenotyper cage was set up for the NOR test. For NOR, one of the objects of the familiarization phase (**Figure S3D**) was replaced with a novel object (parrot, hippo, or giraffe) (**Figure S3F**). The animal was then placed in the cage and was allowed to explore for 5 minutes. After the test, the cumulative duration of the times in which the nose point was in the boundaries of the familiar object and novel object zones and the frequency of entries into each zone was measured using Ethovision software.

#### Novel Location Recognition-24 hours

24 hours post familiarization, the NLR and NOR tests were performed. The procedures used were the same as NLR performed at 30 minutes post familiarization. However, the novel location used in this test was different than the previously set up novel location (**Figure S3G**). After the test, the animal was returned to the home cage and allowed to recover for 5 minutes to prepare for the longterm NOR test.

#### Novel Object Recognition-24 hours

While the animal was recovering in the home cage after the NLR test, the cage was set up for NOR test. Again, the procedures were performed similar to the post-familiarization NOR test; however, the novel object used (elephant) was different from the novel object used last time (**Figure S3H**). After the test, the animals were returned to the home cage and were placed back in the regular housing room.

### Morris Water Maze (MWM) And Reversal Learning

#### Morris Water Maze

~ 2 hours after the end of the NOR and NLR tests, the first phase of MWM was initiated. MWM was done in three phases (Bromley-Brits et al., 2011). Phase 1: Day 0, Flag day. Phase 2: Days 1-4 training. Phase three: Day 5, testing (**Figure S4A**). After the testing day, the reversal learning paradigm was performed (see below). Before initiation of the experiment, all cages in which the animals were singly housed were transferred to the experimental room and covered with a breathable sheet for ~30 minutes, to recover before start of the experiment. All animals were covered the whole time unless being tested. Moreover, during the experiments, the experimenter would be unobservable to the animals to allow for more free behavior of the animal. Also, during all the phases, the animals were released into the pool facing the wall of the pool.

The pool was videotaped during all phases. The pool was divided into 4 quadrants: South-West (SW), South-East (SE), North-East (NE), North-West (NW). To provide visual cues, four different contrasting cues were placed on the pool wall at 90 degrees apart (**Figure S4B**). The water was temperature controlled (~70°C) and was drained and the pool was cleaned with 70% ethanol once every other day.

##### Phase 1

The purpose of phase 1 is to familiarize the animals with the location of the platform in the pool and, as mice do not like to swim, it allows them to identify and learn the location of a place that allows them to stop swimming. For this phase, the pool was filled with water (temperature ~70°C) and the platform was placed in the NE quadrant (Same quadrant where the platform was hidden in the MWM training days). The platform was placed ~1 cm above the water. The water was clear to allow visualization of the platform. The platform was labeled with a red flag for easier observation and finding. One mouse at a time was released gently in the water in the SW quadrant and was allowed to find the platform in 60 seconds. If the animal did not find the platform in 60 seconds, the animal was grabbed by base of the tail and was placed on the platform for 10 seconds.

##### Phase 2 (Training)

The animals were trained for 4 consecutive days for 4 times each day. The trainings were all performed between 2-6 PM. The pool was filled with water (temperature ~70°C). White, non-toxic tempura paint was used to make the water opaque. The platform was placed in the NE quadrant, 1 cm beneath water surface. During each training day, each animal was trained four times to find the platform. Each time, the animal was released into the pool in a different quadrant. The release quadrant order was SW, SE, NE, NW. After the animal was released in the assigned quadrant, it was allowed to investigate and find the platform within 60 seconds and stay on the platform for 5 seconds. If it could not find the platform after 60 seconds, the animal was grabbed by the base of the tail and placed on the platform and let sit for 10 seconds. After each training, the animal was dried with a towel and returned to the home cage and allowed to recover for 15 minutes before the next training on the same day.

##### Phase 3 (Testing)

On the testing day, the platform was removed. During the testing, each animal was placed in the pool at the furthest point from the platform (Middle of SW and SE quadrant) and was allowed to swim for 60 seconds. The total time spent in each individual quadrant was measured.

#### Reversal Morris Water Maze Learning (rMWM)

~24 hours after the MWM testing day, the animals were retrained for two consecutive days, 4 times each day. In the rMWM training, the animals were trained to learn the new location of the platform (SW quadrant). The training was performed similar to the MWM training procedure; however, the platform was not submerged under the water and was 1 cm above the water, visible to the animals.

On the test day, the platform was removed, and the animal was placed in a starting point furthest from the platform location (Between NE and NW). The animal was released into the pool and was allowed to swim for 60 seconds. The total time spent in each quadrant, the total distance traveled, and the velocity of the animal was measured and compared across all groups.

### Statistical inference and data plotting

The tests used for statistical analysis were One-sample Student’s T-test, One-way ANOVA, or Two-way ANOVA. For multiple comparison analysis, Tukey or Sidak tests were used depending on the experimental design. The specific statistical test performed is listed in each corresponding figure legend. In the experiments with n ≤ 12, individual data points were indicated in the graph with mean and standard error of the mean (SEM). The experiments with 12 or more data points were plotted as a bar graph with indication of mean and SEM.

