## Supplementary material for "Spinophilin limits GluN2B-containing NMDAR activity and sequelae associated with excessive hippocampal NMDAR function": All Supplemental Files

**Figure S1. Schematic of protocol and detergents used to isolate crude cytosolic (S1), membrane-associated (S2), and synaptic (S3) fractions.** Hippocampi from P28 WT or spinophilin KO mice were homogenized in an isotonic KCl buffer without detergent and centrifuged at 100,000 x g for 1 hour at 4ºC. The supernatant was mixed with 4X sample buffer and this generated the S1, cytosolic fraction. The pellet was resuspended in the same isotonic KCl buffer containing 1% Triton-X 100. This resuspension was centrifuged at 14,000 x g for 10 minutes at 4ºC and the supernatant was mixed with 4X sample buffer to generate the S2, membrane associated fraction. The pellet was resuspended in the KCl buffer containing 1% Triton-X 100 and 1% deoxycholate and sonicated. This was mixed with 4X sample buffer to generate the S3, synaptic fraction.

**Figure S2. Schematic of neuronal stress protocol on hippocampal neurons.** Primary hippocampal cultures were assayed at 14-24 DIV. On the test day, the culture media was collected and set aside as conditioned Neurobasal (cNB) media. 1 mL of fresh neurobasal media alone or containing 100 µM glutamate or glutamate + AP-5 was added to the wells. The cells were incubated with the media for 30 minutes. After 30 minutes, the media was removed and replaced with 1.5 mL cNB media for 90 minutes.

**Figure S3. Location and objects used for the novel location and novel object tests.** Four different objects were used for the novel location and novel object tests (S3A). Animals were habituated to the Phenotyper cage daily for 3 days (S3B). Zones were set up around the different objects (S3C) and the initial day had two of the same objects placed in the arena (S3D). Mice were allowed to explore the objects for 20 minutes then were returned to their home cage. 30-minutes following the test, mice were placed back in the Phenotyper cage with one of the objects moved from the top right to bottom right quadrant (S3E). The mouse was allowed to explore for 5 minutes and the time interacting with each object was recorded. The mouse was removed back to its home cage for 5 minutes and the novel location object was removed and a novel object was placed in the top-right quadrant (S3F). The time exploring the objects was recorded for 5 minutes. The novel location test was repeated 24 hours later with a new location (bottom left quadrant, S3G) and the novel object test was repeated with a $2^{nd}$ novel object (S3H).

**Figure S4. Schematic and outline for the Morris Water Maze (MWM) and reversal MWM (rMWM) training.** On day 0, mice were placed in between the SW and SE zones of the maze that was filled with clear water. The platform was located 1-cm below the water in the NE zone and was marked with a red flag. On training days 1-4, the water was colored white with tempura paint and the mice were released from in between each of the four zones SW/SE, SE/NE, NE/NW, NW/SW. The time to find the platform was recorded and the mouse was allowed to sit on the platform for 5 seconds. If the mouse did not find the platform, it was placed on the platform for 10 seconds. On day 5 the platform was removed and the mice were released from in between the SW/SE zones. The time spent in each zone was recorded. For the rMWM, the platform position was moved and the platform was placed in the SW quadrant, using a visible (1-cm above the water) platform. The mice were trained in this way for 2 days and then on the $3^{rd}$ day, the platform was removed.

**Figure S5. Latency and percentage of animals that found the platform during the MWM and rMWM. A.** Latency to find the hidden platform on training days 1-4 in MWM test. **B.** Percentage of animals that found the submerged platform within 60 seconds in MWM test. **C.** Latency to find the visible platform on reversal training days 1-2 in rMWM test. **D.** Percentage of animals that found the visible platform within 60 seconds in rMWM test.

Figure S1

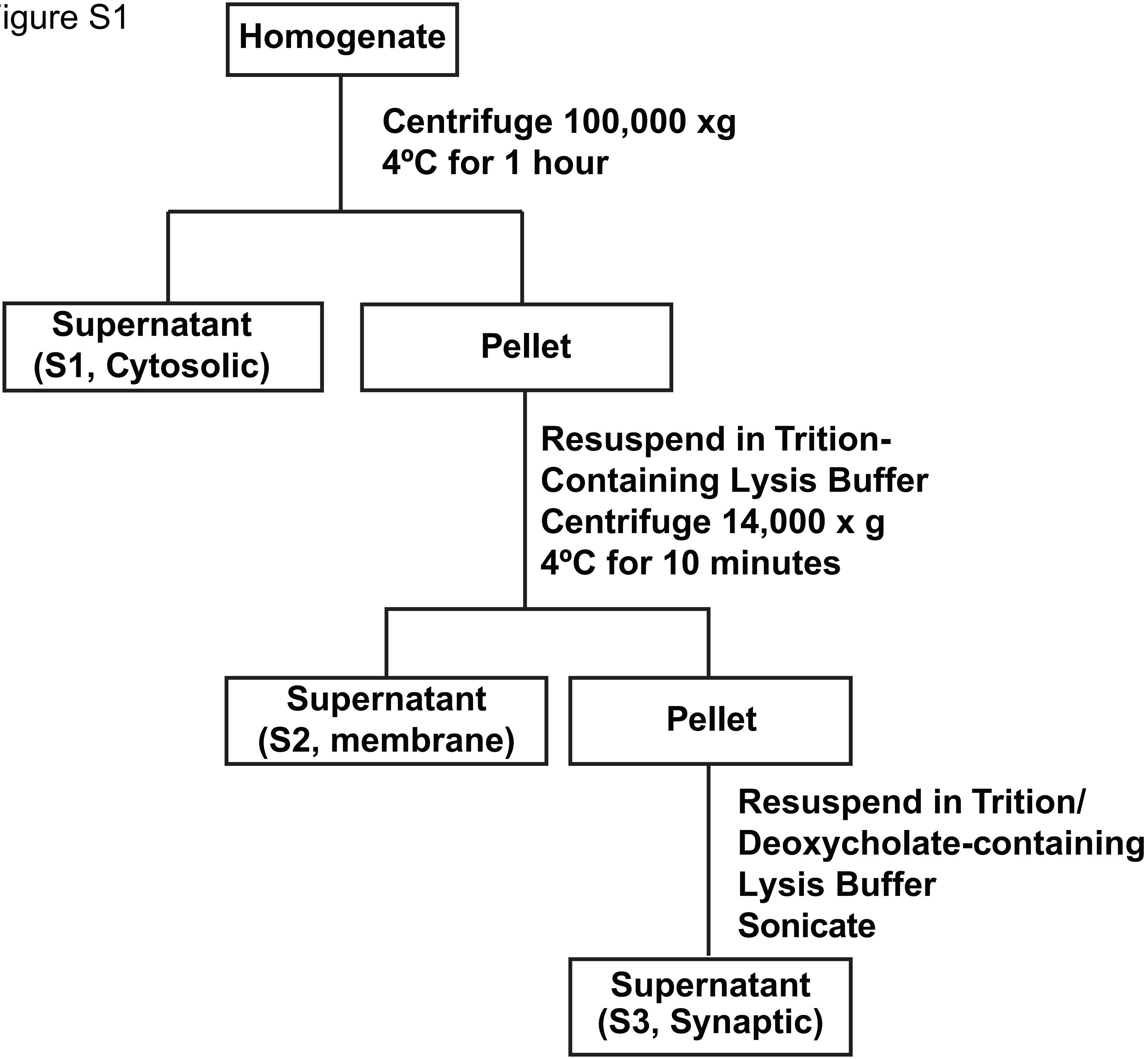
Homogenate
Centrifuge 100,000 xg
4ºC for 1 hour
Supernatant
(S1, Cytosolic)
Pellet
Resuspend in Trition-
Containing Lysis Buffer
Centrifuge 14,000 x g
4ºC for 10 minutes
Supernatant
(S2, membrane)
Pellet
Resuspend in Trition/
Deoxycholate-containing
Lysis Buffer
Sonicate
Supernatant
(S3, Synaptic)

Figure S2

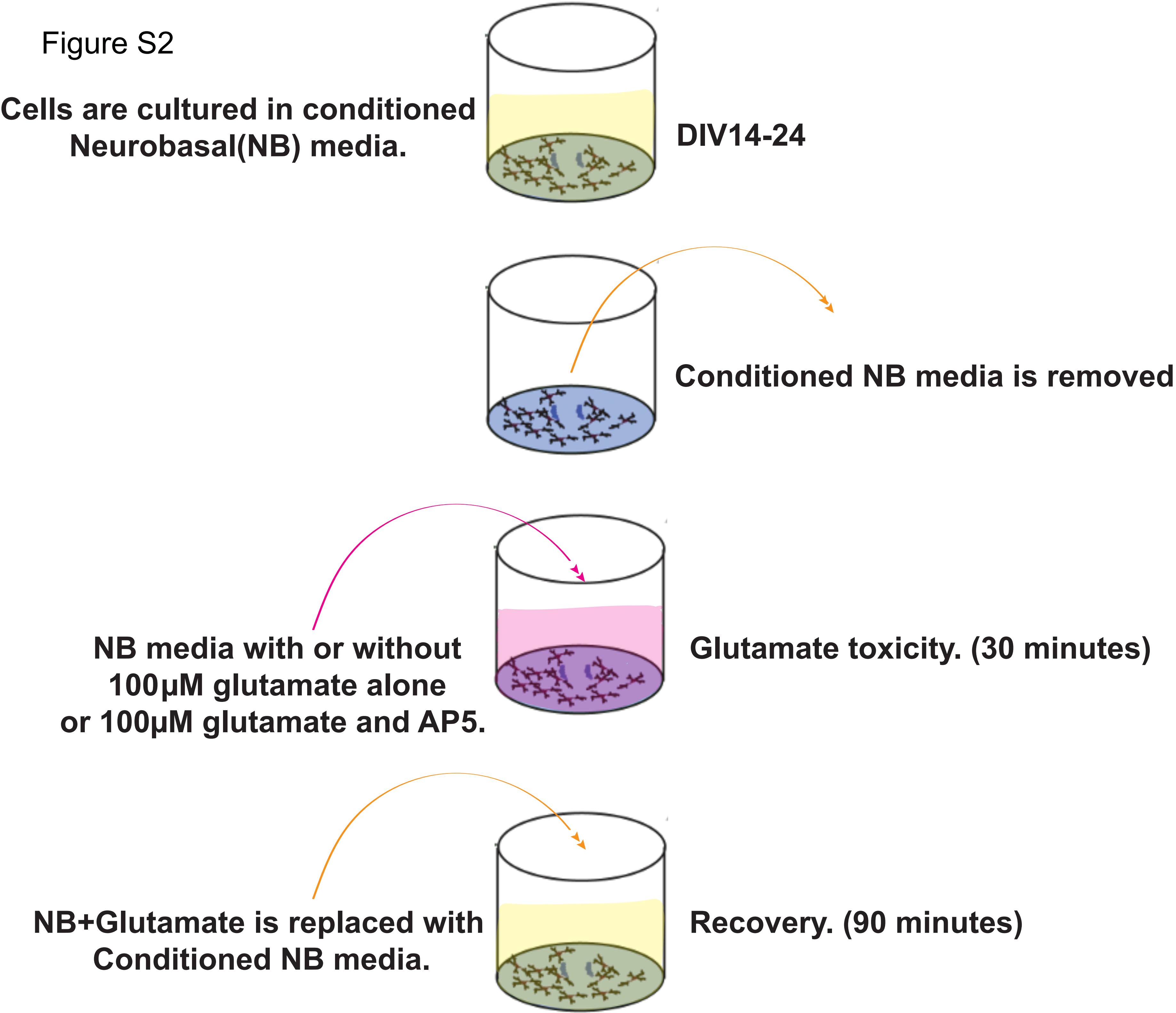
Cells are cultured in conditioned Neurobasal(NB) media.
DIV14-24
Conditioned NB media is removed
NB media with or without 100µM glutamate alone or 100µM glutamate and AP5.
Glutamate toxicity. (30 minutes)
NB+Glutamate is replaced with Conditioned NB media.
Recovery. (90 minutes)

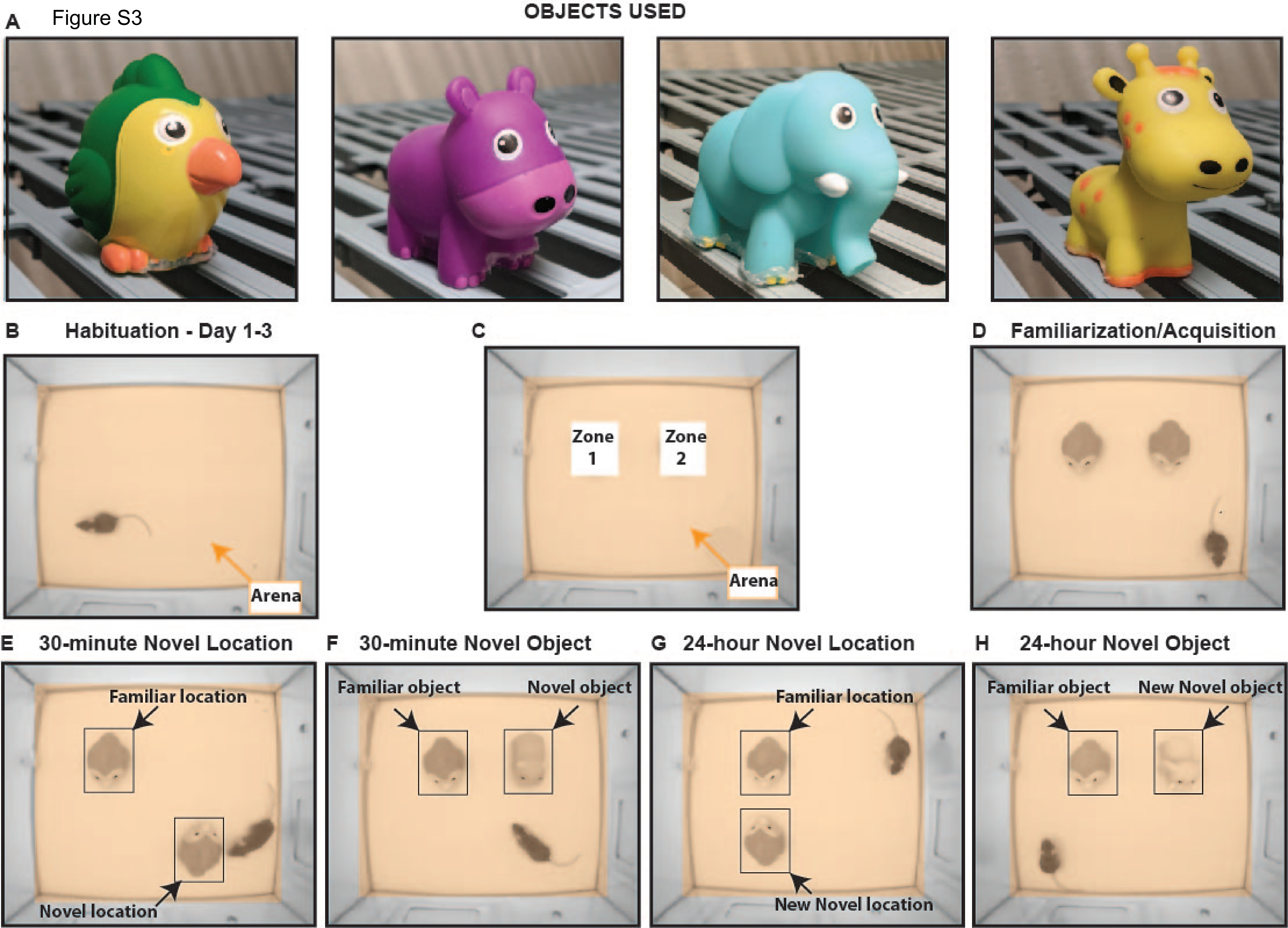

A
Figure S3
OBJECTS USED
B
Habituation - Day 1-3
Arena
C
Zone 1
Zone 2
Arena
D
Familiarization/Acquisition
E
30-minute Novel Location
Familiar location
Novel location
F
30-minute Novel Object
Familiar object
Novel object
G
24-hour Novel Location
Familiar location
New Novel location
H
24-hour Novel Object
Familiar object
New Novel object

Figure S4

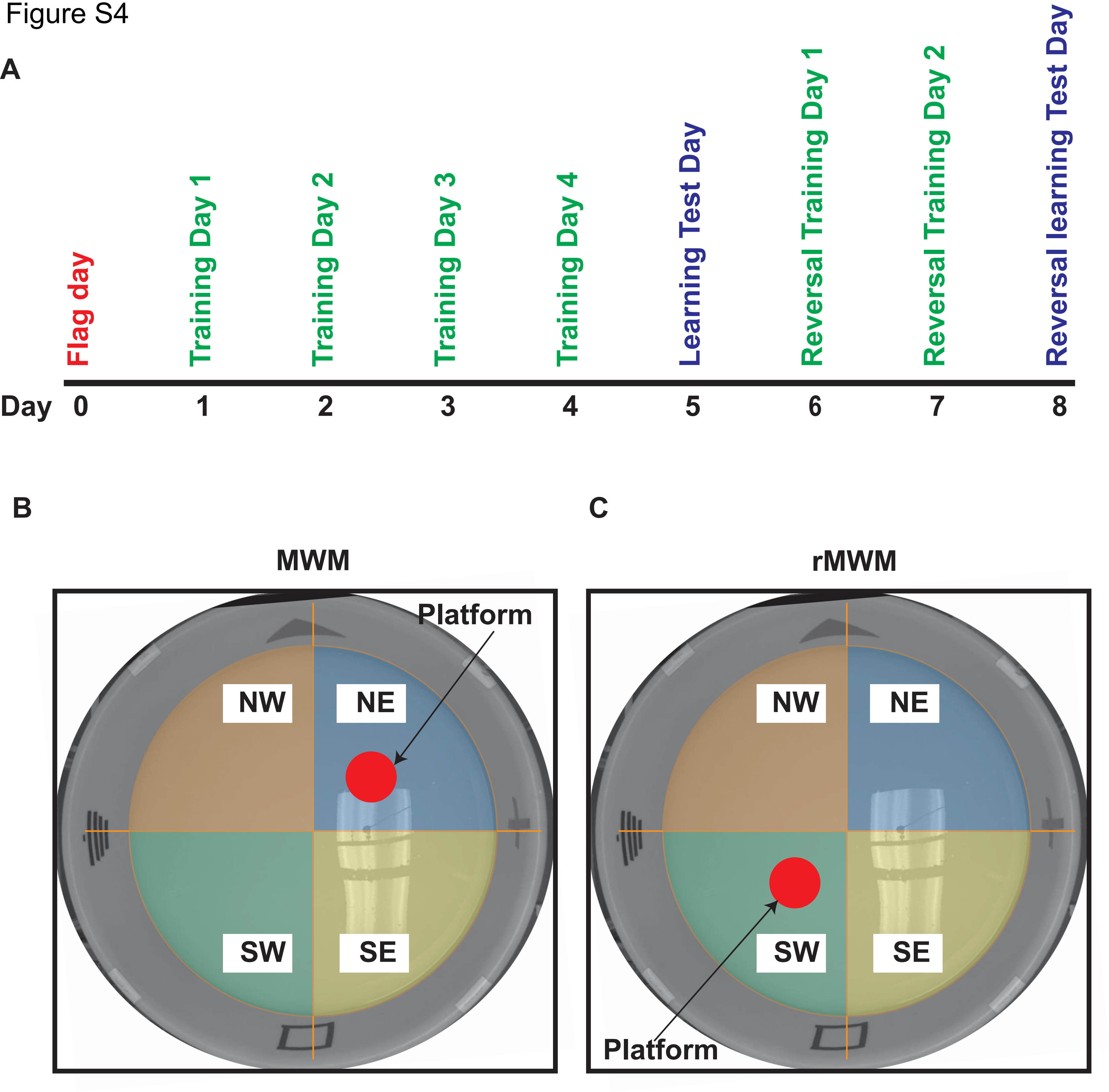
A
Flag day
Training Day 1
Training Day 2
Training Day 3
Training Day 4
Learning Test Day
Reversal Training Day 1
Reversal Training Day 2
Reversal learning Test Day
Day 0 1 2 3 4 5 6 7 8
B
MWM
Platform
NW
NE
SW
SE
C
rMWM
NW
NE
SW
SE
Platform

Figure S5

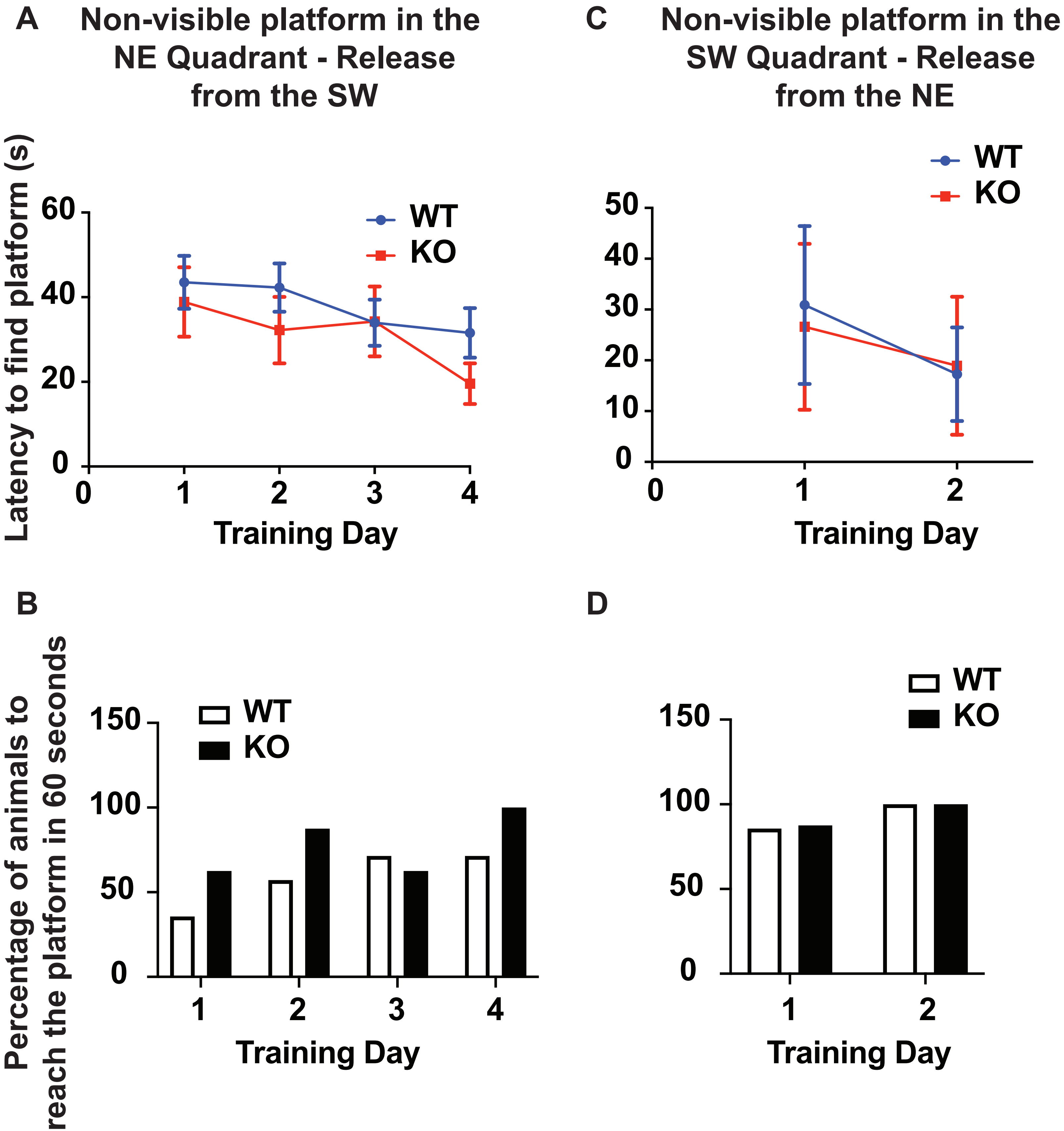
A
Non-visible platform in the NE Quadrant - Release from the SW
Latency to find platform (s)
60
40
20
0
0
1
2
3
4
Training Day
WT
KO
C
Non-visible platform in the SW Quadrant - Release from the NE
50
40
30
20
10
0
0
1
2
Training Day
WT
KO
B
Percentage of animals to reach the platform in 60 seconds
150
100
50
0
1
2
3
4
Training Day
WT
KO
D
150
100
50
0
1
2
Training Day
WT
KO
